# Eugenol mimics exercise to promote skeletal muscle fiber remodeling and myokine IL-15 expression by activating TRPV1 channel

**DOI:** 10.1101/2023.07.18.549410

**Authors:** Tengteng Huang, Xiaoling Chen, Jun He, Ping Zheng, Yuheng Luo, Aimin Wu, Hui Yan, Bing Yu, Daiwen Chen, Zhiqing Huang

## Abstract

Metabolic disorders are highly prevalent in modern society. Exercise mimetics are defined as pharmacologic compounds that can produce the beneficial effects of fitness. Recently, there has been increased interest in the role of eugenol and transient receptor potential vanilloid 1 (TRPV1) in improving metabolic health. The aim of this study was to investigate whether eugenol acts as an exercise mimetic by activating TRPV1. Here, we showed that eugenol improved endurance capacity, caused the conversion of fast to slow muscle fibers, and promoted white fat browning and lipolysis in mice. Mechanistically, eugenol promoted muscle fiber type transformation by activating TRPV1-mediated CaN signaling pathway. Subsequently, we identified IL-15 as a myokine that is regulated by the CaN/Nuclear factor of activated T cells cytoplasmic 1 (NFATc1) signaling pathway. Moreover, we found that TRPV1-mediated CaN/NFATc1 signaling, activated by eugenol, controlled IL-15 levels in C2C12 myotubes. Our results suggest that eugenol may act as an exercise mimetic to improve metabolic health via activating the TRPV1-mediated CaN signaling pathway.

## Introduction

Due to the lack of regular exercise among many individuals (***Carlson et al., 2010***), metabolic diseases are highly prevalent in modern society (***Hoehn et al., 2010***). To address this issue, pharmacological interventions have been considered as potential treatments. Exercise mimetics are pharmacologic compounds that produce fitness benefits (***Fan and Evans, 2017***). Although some synthetic exercise mimetics such as AICAR (AMPK activator), GW501516 (PPARδ activator), SRT1720 (SIRT1 activator), and GSK4716 (ERRγ activator) have been used to improve fitness (***Feige et al., 2008; Narkar et al., 2008; Rangwala et al., 2010***), the safety and health of each drug must be considered. In recent years, plant extracts such as resveratrol, lycium barbarum and epicatechin have demonstrated potential as a new class of low-toxin exercise mimetics (***Meng et al., 2020; Nogueira et al., 2011; Wen et al., 2020***). Therefore, it is of great significance to discover natural medicinal and edible plants that have mimetic effects of exercise.

Based on their metabolism and contractile properties, muscle fibers are commonly divided into slow oxidative fibers (muscle fibers that express myosin heavy chain (MyHC) I) and fast glycolytic/oxidative fibers (muscle fibers that express MyHC IIa, MyHC IIx, or MyHC IIb) (***Schiaffino and Reggiani, 2011***). Different muscle fibers exhibit a high degree of plasticity and are regulated by exercise (***Egan and Zierath, 2013***). The increased proportion of slow muscle fibers contributes to the enhancement of mitochondrial biogenesis and lipid metabolism (***Carlson et al., 2010***). Therefore, one of the benefits of exercise is to treat metabolic disorders by promoting slow muscle fiber remodeling (***Duan et al., 2017***). In addition, it has been increasingly recognized that myokines released by skeletal muscle play an essential role in mediating the benefits of exercise. Myokines are defined as cytokines or peptides that are released from skeletal muscle cells, which exert autocrine, paracrine, or endocrine effects (***Pedersen and Febbraio, 2012***). As a medium of crosstalk between muscles and other organs, myokines exert systemic regulation of exercise by acting on the muscle, fat, liver, pancreas, and other organs, effectively improving insulin resistance, obesity, and metabolic disorders of type 2 diabetes (***Severinsen and Pedersen, 2020***). Therefore, another potential effect of exercise is to induce the release of myokines (***Fan and Evans, 2017***). Although many previous studies have discovered plant extracts as exercise mimetics by promoting slow oxidative muscle fiber, few studies have investigated the effect of exercise mimetics on the release of myokines.

Eugenol is a safe natural compound extracted from clove oil and various plant spices such as basil, bay leaves, and cinnamon (***Kaur et al., 2010***). Eugenol exhibits a variety of biological activities, including antioxidative (***Magalhães et al., 2019***), antibacterial (***Devi et al., 2010***), anti-inflammatory (***Magalhães et al., 2019***) and anti-cancer activities (***Lesgards et al., 2014***). Recent studies have also suggested that eugenol could be a promising therapeutic drug for preventing diabetes and obesity (***Al-Trad et al., 2019; Jung et al., 2012; Mnafgui et al., 2013; Srinivasan et al., 2014***), indicating its potential as an exercise mimetic. In addition. as a member of the transient receptor potential (TRP) channel family, TRP vanilloid 1 (TRPV1), also known as the capsaicin receptor, has been reported to improve endurance capacity and energy metabolism (***Luo et al., 2012***), counter obesity (***Baskaran et al., 2016***), and intervene diabetes (***Wang et al., 2012***), making it a potential target protein for discovering exercise mimetics.

Eugenol contains a vanilloyl fragment that may bind to TRPV1 similarly to capsaicin, implying that eugenol might exert its biological activity via TRPV1 (***Harb et al., 2019***). It has been found that eugenol activates TRPV1 in a heterologous expression system and rat trigeminal ganglion neurons (***Xu et al., 2006; Yang et al., 2003***). Furthermore, a recent study demonstrated that eugenol activated TRPV1 in skeletal muscle (***Jiang et al., 2022***). TRPV1 is an important Ca^2+^ entry pathway contributing to increase intracellular Ca^2+^ (***Nilius et al., 2007***), suggesting that TRPV1 may regulate many biological processes through Ca^2+^-dependent signaling pathways. As a Ca^2+^-dependent signaling, calcineurin (CaN) is a core signaling to promote fast to slow muscle fiber transformation (***Sakuma and Yamaguchi, 2010***). Additionally, CaN signaling plays an essential role in regulating the expression of myokine IL-6 (***Banzet et al., 2005; Banzet et al., 2007***). Therefore, we hypothesize that eugenol, as an exercise mimetic, promotes the release of myokines and fast to slow muscle fiber transformation via the TRPV1-mediated CaN signaling pathway. The primary objective of this study is to investigate this hypothesis.

## Results

### Eugenol promotes fast to slow muscle fiber transformation in mice and in C2C12 myotubes

As depicted in Figure 1A and B, eugenol did not have any effect on the body weight or skeletal muscle weight in mice. However, the skeletal muscle in the EUG50 and EUG100 groups exhibited a redder muscle color (Figure 1C), indicating a shift from fast to slow muscle fiber in these two groups. Further experiments showed that EUG50 and EUG100 increased the expression of slow MyHC and decreased the expression of fast MyHC protein in GAS and TA muscles (Figure 1D and E). Additionally, the mRNA expression of *MyHC I*, *MyHC IIa*, *MyHC IIx*, and *MyHC IIb* was generally consistent with protein expression (Figure 1G). As our *in vivo* studies demonstrated that the EUG200 group had no effect on muscle fiber type, we suspected that high doses of eugenol may have no impact on muscle fiber type. Therefore, we selected a broad range of eugenol doses (0-200 μM) based on the safe range of eugenol doses determined by the CCK-8 assay (Figure 1-figure supplement 1) to treat C2C12 myotubes and replicate these results *in vivo*. As indicated in Figure 1F, 12.5-50 μM eugenol boosted slow MyHC expression, while 12.5-100 μM eugenol decreased fast MyHC expression. Furthermore, as shown in Figure 1H, 100 μM eugenol increased the mRNA expression of *MyHC I*, and 12.5-100 μM eugenol increased the mRNA expression of *MyHC IIa*, whereas 50 μM eugenol decreased the mRNA expression of *MyHC IIb*. Consistent with our in vivo studies, our in vitro findings again suggested that high doses (200 μM) of eugenol had no effect on muscle fiber type. In summary, our results suggest that eugenol promotes a transformation from fast to slow muscle fiber. However, it should be noted that high doses of eugenol may have no effect on muscle fiber type.

**Figure 1.**
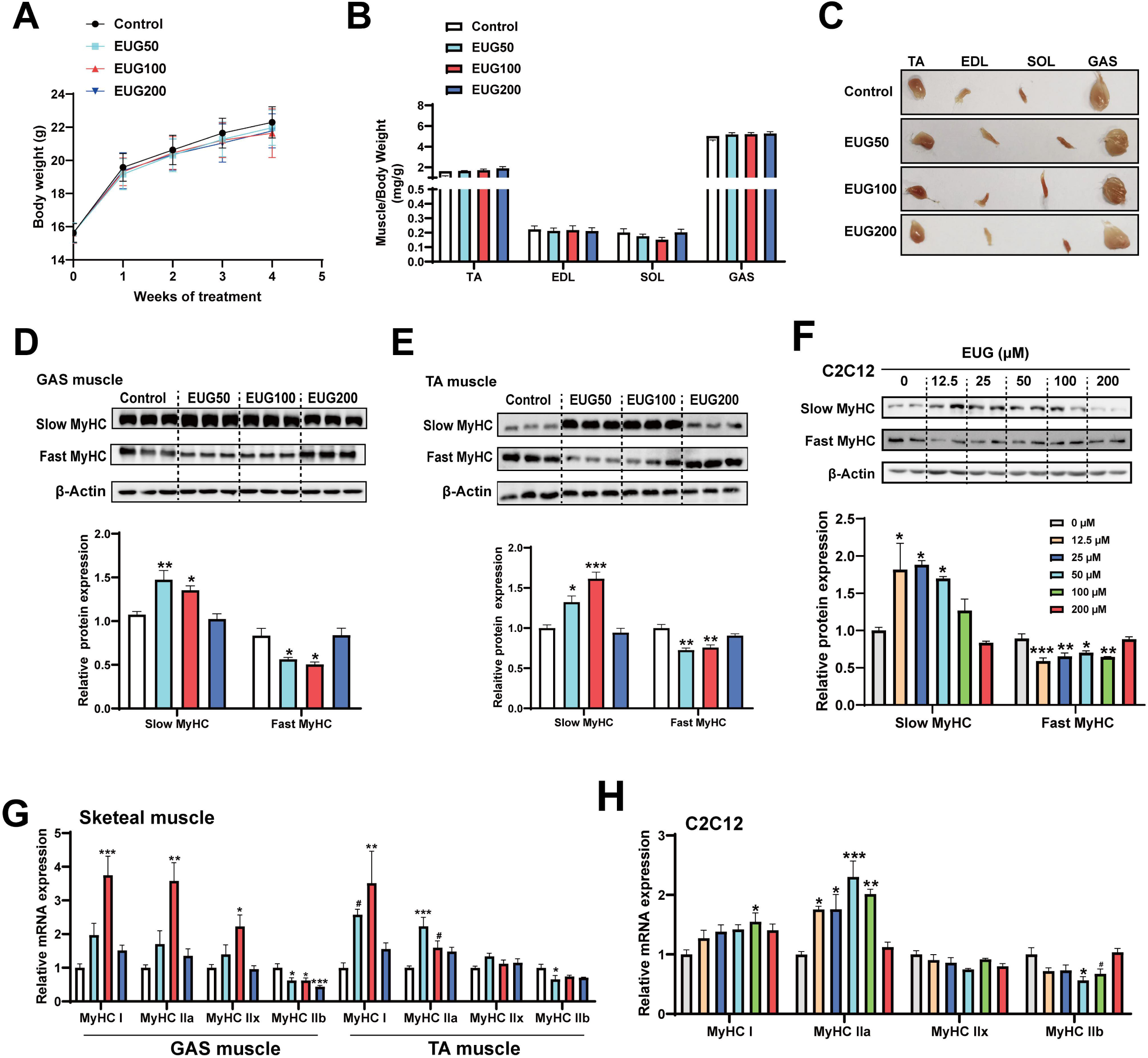
Eugenol promotes the transformation of fast to slow musle fiber. **(A)** The body weight of the mice. (**B, C)**, Skeletal muscle weight and representative images of skeletal muscle. **(D-F)** The protein expression of muscle fiber type in GAS and TA muscle and in C2C12 myotubes. (**G-H)** The mRNA expression of muscle fiber type in GAS and TA muscle and in C2C12 myotubes. For A-B, N=12 per group. For D-E, N=3 per group. For F and H, N=4 per group. For G, N=6 per group. \**P* < 0.05, \*\**P* < 0.01, and \*\*\**P* < 0.001.

### Eugenol promotes oxidative metabolism activity, mitochondrial function, and endurance performance

An increase in the proportion of slow muscle fibers is often accompanied by an increase in skeletal muscle oxidative metabolism activity, mitochondrial function, and endurance performance. Therefore, we examined these indicators in our study. As shown in Figure 2A, EUG100 increased the exhaustion time of mice. As shown in Figure 2B, EUG50 and EUG100 decreased LDH activity and increased SDH activity in the GAS muscle. In the TA muscle, LDH activity was decreased in all EUG groups, and EUG100 increased the activities of SDH and MDH. Then, we chose 100 mg/kg eugenol (as the optimal dose) for the exhausting swimming test. In addition, the transcript levels of mitochondrial transcription factors *PGC-1α*, *NRF1*, and *TFAM* and the mRNA expression of components of the mitochondrial electron transport chain were increased in eugenol-treated mice (Figure 2C). The protein expression of mitochondrial electron transport complex I and complex V was upregulated in eugenol-treated mice (Figure 2D). Furthermore, EUG100 improved the mtDNA copy number in the GAS and TA muscle (Figure 2E), and 12.5-100 μM EUG increased the mtDNA copy number in C2C12 myotubes (Figure 2F).

**Figure 2.**
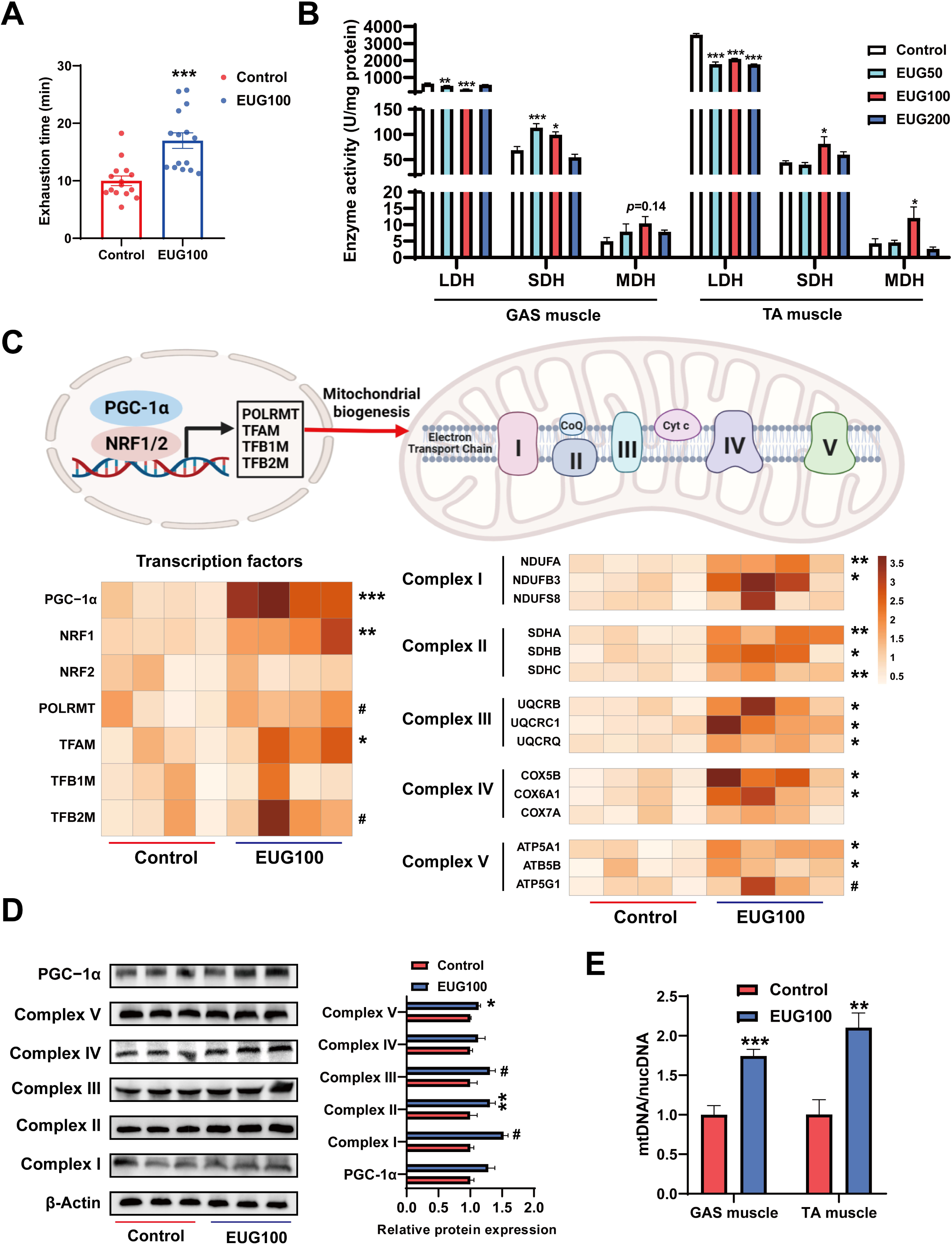
Eugenol promotes oxidative metabolism activity, mitochondrial function, and endurance performance of mice. **(A)** The effect of eugenol on exhausting swimming time in mice. **(B)** The effect of eugenol on metabolism enzymes activity in GAS and TA muscle. (**C)** The heatmap for the mRNA expression of genes encoding mitochondrial complex components and transcription factors controling mitochondrial biogenesis in GAS muscle. Color gradient represents relative mRNA expression with darker colors indicating higher expression. **(D)** Protein expression of mitochondrial electron transport complexes in GAS muscle. Complex I (NDUFA1), complex II (SDHA), complex III (UQCRC1), complex IV (MTCO1), and comlex V (ATP5B). (**E)** mtDNA copy number in muscles and cells. For A, N=15 per group. For B, N=6 per group. For C and F, N=4 per group. For D, N=3 per group. For E, N=6 per group. ^#^*P*<0.1, \**P* < 0.05, \*\**P* < 0.01, and \*\*\**P* < 0.001.

### Eugenol promotes lipolysis and browning of fat

Our research has found that both EUG100 and EUG200 promoted average daily feed intake (ADF) (Figure 3A). However, interestingly, there was no change in body weight (Figure 1A). This indicates that there was an increase in ADF/average daily weight gain (ADG) (Figure 3A). In addition, eugenol decreased iWAT and gWAT weight while promoting BAT weight (Figure 3B). And eugenol decreased T-CHO and LDL while increasing HDL level in serum (Figure 3C). These apparent results suggest that eugenol may increase the energy metabolism rate in mice, offsetting the weight gain and increased fat synthesis resulting from increased food intake. Therefore, we further speculate that eugenol may promote lipolysis and fat thermogenesis. As we speculated, we found that eugenol promoted the mRNA expression of the fat synthesis-related genes *PPARγ* and *HSL*, as well as the fatty acid transport gene *FABP4*, in iWAT (Figure 3D). In gWAT, it was found that eugenol promoted the mRNA expression of the *FASN* and *FABP4* (Figure 3D). We further examined the mRNA expression of key genes involved in browning of fat, and found that eugenol promoted the mRNA expression of *CD137*, *TBX1*, *UCP-1*, *PRDM16*, *Dio2*, and *Cidea* in iWAT, while eugenol promoted the mRNA expression of *TMEM26*, *UCP-1*, *PRDM16*, *Dio2*, and *Cidea* in gWAT (Figure 3E). At the protein level, eugenol promoted the expression of the fatty acid transport protein FABP-1, as well as the UCP-1 protein expression in gWAT (Figure 3F). In addition, we also examined the effects of eugenol on the browning-related proteins and mitochondrial complex proteins in BAT. It was found that eugenol promoted the UCP-1 and PGC-1α protein expression (Figure 3G), as well as the mitochondrial electron transport complex III and complex V protein expression (Figure 3H). Together, these results indicate that eugenol promotes lipolysis and browning of fat.

**Figure 3.**
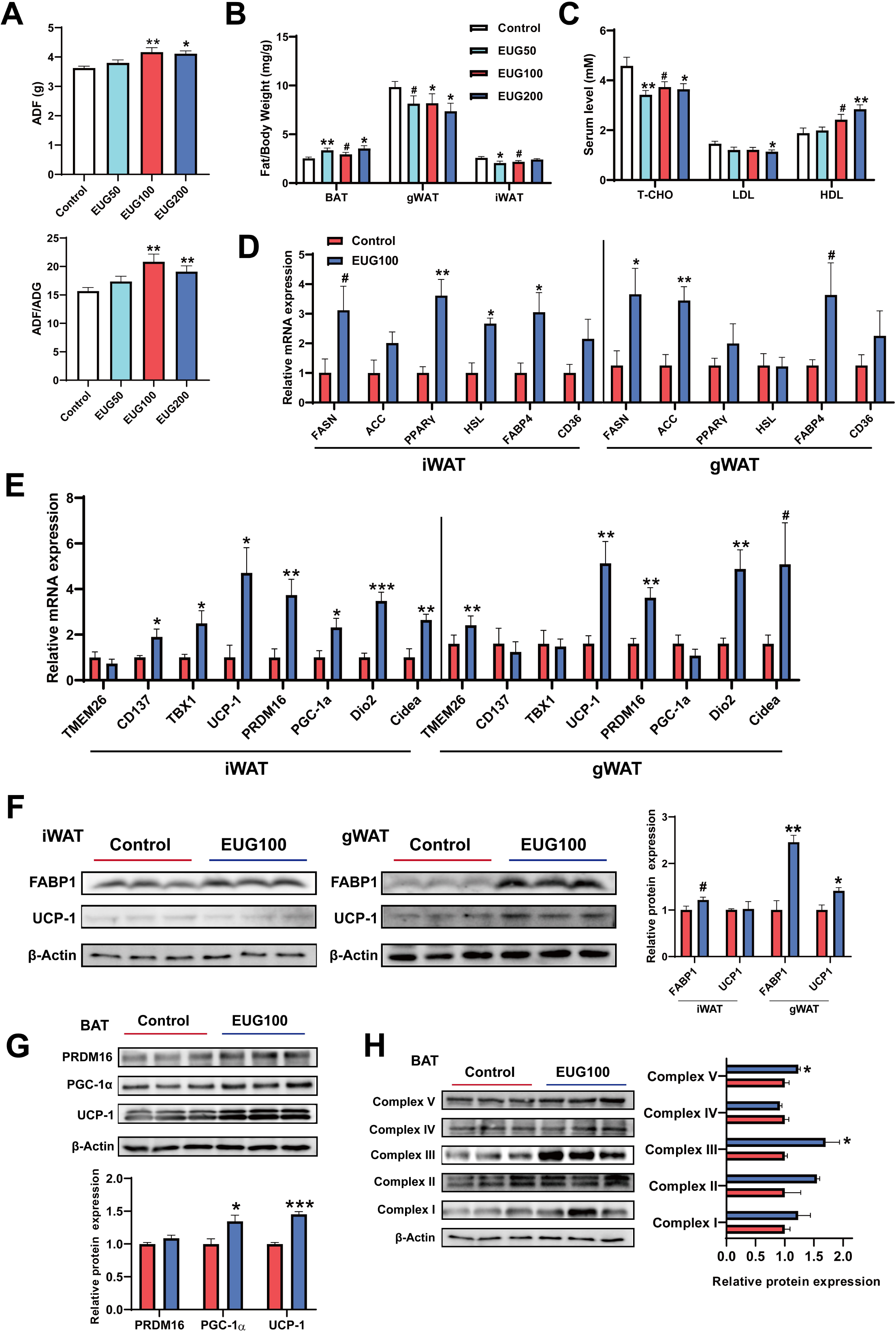
Eugenol enhances lipolysis and thermogenesis. (**A)** The ADF and the ration of ADF to ADG. (**B)** Tissue weight of adipose weight. (**C)** The level of T-CHO, LDL, and HDL in serum. (**D)** The mRNA expression of genes related to lipolysis, lipogenesis, and lipid transport in iWAT and gWAT. (**E)** The mRNA expression of genes related to adipose browning and thermogenesis in iWAT and gWAT. (**F)** The protein expression of FABP1 and UCP1 in iWAT and gWAT. (**G)** The expression of protein related to adipose browning and thermogenesis in BAT. (**H)** The protein expression of mitochondrial electron transport complexes in BAT. For A-B, N=12 per group. For C-E, N=6 per group. For F-H, N=3 per group. ^#^*P*<0.1, \**P* < 0.05, \*\**P* < 0.01, and \*\*\**P* < 0.001.

### Eugenol activates TRPV1-mediated CaN/NFATc1 signaling pathway in skeletal muscle

Based on the gene expression profiling, it was observed that TRPV1 is expressed in all tissues, including adipose and muscle tissues, with the highest expression in skeletal muscle compared to other TRP channels (Figure 4A and Figure 4-figure supplement 1A). qPCR analysis confirmed the expression of TRPV1 and TRPV2 in skeletal muscle, and the mRNA expression of only *TRPV1* was promoted by EUG50 and EUG100 in TA muscle. In C2C12 cells, only TRPV1-4 were expressed, and only *TRPV1* mRNA expression was promoted by 25 and 50 μM eugenol (Figure 4C). Adipose tissue expressed TRPV1 and TRPV2 genes, and EUG100 and EUG200 promoted *TRPV1* mRNA expression, while EUG50 promoted T*RPV2* mRNA expression (Figure 4-figure supplement 1B and C). The effect of eugenol on TRPV1 protein expression was consistent with its effect on *TRPV1* mRNA expression in both skeletal muscle tissue and C2C12 cells (Figure 4D-F). Moreover, taking the TRPV1-capsaicin binding sites (TYR511, SER512, THR550, and GLU570) (***Carnevale and Rohacs, 2016***) as the potential binding pocket, molecular docking analysis showed that eugenol bound at the binding pocket and interacted with THR550, ASN551, LEU553, TYR554, ALA566, ILE569, GLU570, and ILE573 (Figure 4-figure supplement 2). Based on the above results, we conclude that eugenol has the potential to activate TRPV1 in both skeletal muscle and adipose tissue. We further investigated the TRPV1-mediated CaN/NFATc1 signaling pathway in skeletal muscle. CnA is a catalytic subunit of CaN, and its expression level reflects the activity of CaN. Our results showed that EUG50 and EUG100 increased CnA protein expression in the GAS and TA muscles (Figure 4D and 4E), and 12.5-100 μM eugenol promoted CnA protein expression in C2C12 myotubes (Figure 4F). The regulator of calcineurin 1 (MCIP1) is a biomarker to reflect the CaN activity (***Yang et al., 2000***), the mRNA expression of *MCIP1* was increased in EUG100 groups (Figure 4-figure supplement 3). Furthermore, the nuclear translocation of NFATc1 was promoted by EUG50 and EUG100 in the GAS and TA muscles, and by 12.5-50 μM eugenol in C2C12 myotubes (Figure 4G-I). In summary, our *in vivo* and *in vitro* studies both suggest that eugenol can activate the TRPV1-mediated CaN/NFATc1 signaling pathway in skeletal muscle. Interestingly, similar to the effects of eugenol on muscle fiber types, we again found that high doses of eugenol have no effect on the signaling pathway.

**Figure 4.**
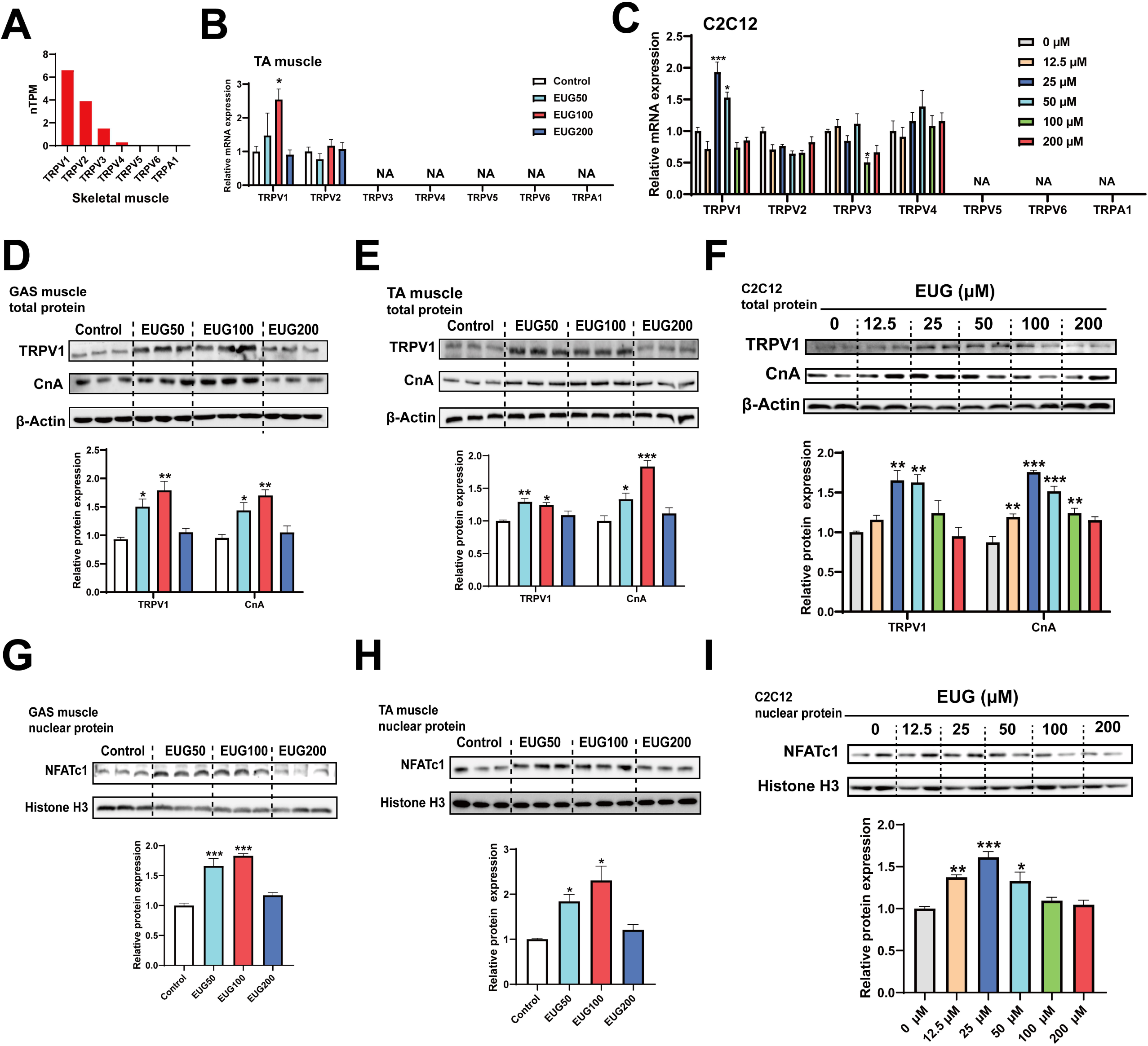
Eugenol activated TRPV1-mediated CaN/NFATc1 signaling pathway. (**A)** The gene expression profile of TRP channels in skeletal muscle was obtained from the GTEx dataset in The Human Protein Atlas (https://www.proteinatlas.org/). (**B, C)** The mRNA expression of TRP channels in TA muscle and C2C12 myotubes. (**D-F)** The TRPV1 and CnA protein expression in GAS and TA muscle and in C2C12 myotubes. (**G-I)** The protein expression of NFATc1 in GAS and TA muscle and in C2C12 myotubes. For B, N=6 per group. For C, F, and I, N=4 per group. For D-E and G-H, N=3 per group. \**P* < 0.05, \*\**P* < 0.01, and \*\*\**P* < 0.001.

### Eugenol promotes fast to slow muscle fiber transformation by activating TRPV1-mediated CaN/NFATc1 signaling pathway

To further investigate the role of TRPV1 in regulating fast to slow muscle fibers, we treated C2C12 myotubes with 25 μM eugenol and either 1 μM TRPV1 inhibitor AMG-517 or 0.5 μM CaN inhibitor cyclosporine A (CsA). The results showed that TRPV1 inhibition weakened the increase in intracellular Ca^2+^ levels induced by eugenol, suggesting that eugenol acts via TRPV1 (Figure 5A). Furthermore, the inhibition of TRPV1 and CaN attenuated the effect of eugenol on CaN (Figure 5B). In addition, eugenol increased the mitochondrial electron transport complex I, II, III, and V protein expression, the inhibition of TRPV1 and CaN attenuated the effect of eugenol on complex I, III, and V (Figure 5C). Importantly, immunofluorescence and Western blot analysis revealed that eugenol promoted the fast to slow muscle fiber transformation, while the inhibition of TRPV1 and CaN eliminated this effect (Figure 5D and Figure 5E).

**Figure 5.**
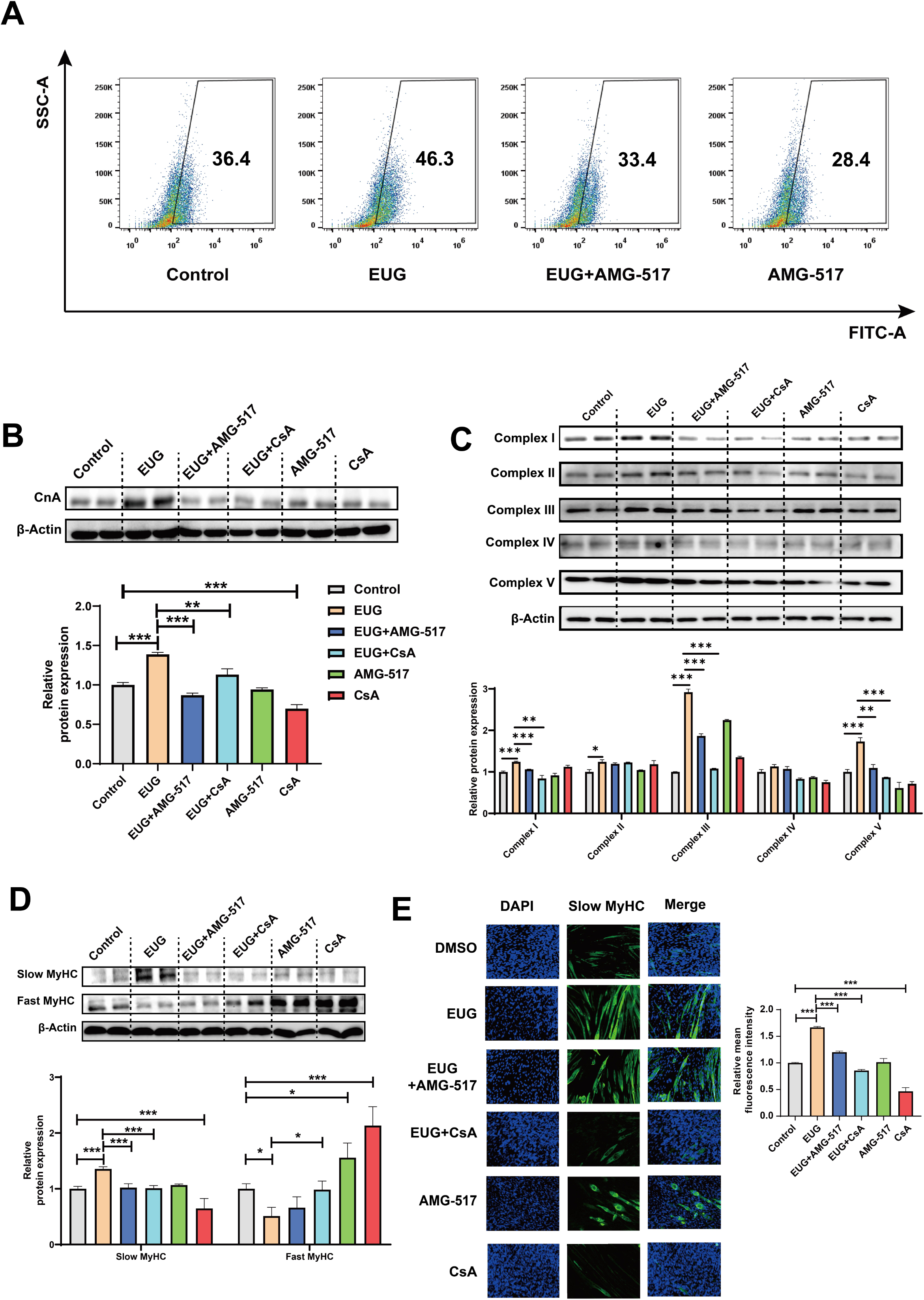
Eugenol promotes fast to slow muscle fiber transformation by activating TRPV1-mediated CaN/NFATc1 signaling pathway. C2C12 myotubes were treated by 25 μM eugenol and 1 μM TRPV1 inhibitor AMG-517 or 0.5 μM CaN inhibitor CsA for 1 day after 4 days of differentiation. (**A)** The flow cytometry assay was used to detect Ca^2+^ levels in C2C12 myotubes; FITC means the fluo-4 fluorescence and SSC means side scatter. (**B)** Western blot was uesd to detect CnA protein expression in C2C12 myotubes. **(C)** Western blot was uesd to detect mitochondrial electron transport complexes protein expression in C2C12 myotubes. (**D)** Western blot was uesd to detect slow MyHC and fast MyHC protein expression in C2C12 myotubes. (**E)** Representative immunofluorescence images of Slow MyHC (Green fluorescence) and relative mean fluorescence intensity quantification. Magnification: × 200. For A-D, N=4 per group. \**P* < 0.05, \*\**P* < 0.01, and \*\*\**P* < 0.001.

### The myokines regulated by CaN

To investigate which myokines are controlled by CaN, C2C12 myotubes were treated with the Ca^2+^ ionophore A23187. As shown in Figure 6A-C, A23187 increased the mRNA expression of *MCIP1*, the protein expression of CnA, and CaN activity. We chose 0.5 μM A23187 for subsequent experiments and confirmed that it increased intracellular Ca^2+^ levels (Figure 6D). To identify CaN-controlled myokines, we next detected the mRNA expression of several myokines that have been well documented to improve metabolic homeostasis (***Eckel, 2019; Whitham and Febbraio, 2016***) and promote fast to slow muscle fiber transformation (***Correia et al., 2021; Knudsen et al., 2020; Men et al., 2021; Quinn et al., 2013***) (Table 1). As shown in Figure 6E, A23187 significantly increased the mRNA expression of *FNDC5*, *IL-6*, *IL-15*, and *Metrnl*. However, *IL-13* mRNA expression was not detected in C2C12 myotubes. In addition, a correlation heatmap showed the highest correlation coefficient (R=0.954) between MCIP1 expression and IL-15 expression, followed by MCIP1 and IL-6 (R=0.846), MCIP1 and Metrnl (0.77), and MCIP1 and FNDC5 (0.651) (Figure 6F). Furthermore, a TFBS prediction revealed more potential binding sites between the IL-15 promoter and NFATc1 (Figure 6G). Based on the correlation and TFBS analysis, it was suggested that IL-15 is the myokine most likely to be regulated by the CaN/NFATc1 signaling pathway, and we selected it for further study.

**Figure 6.**
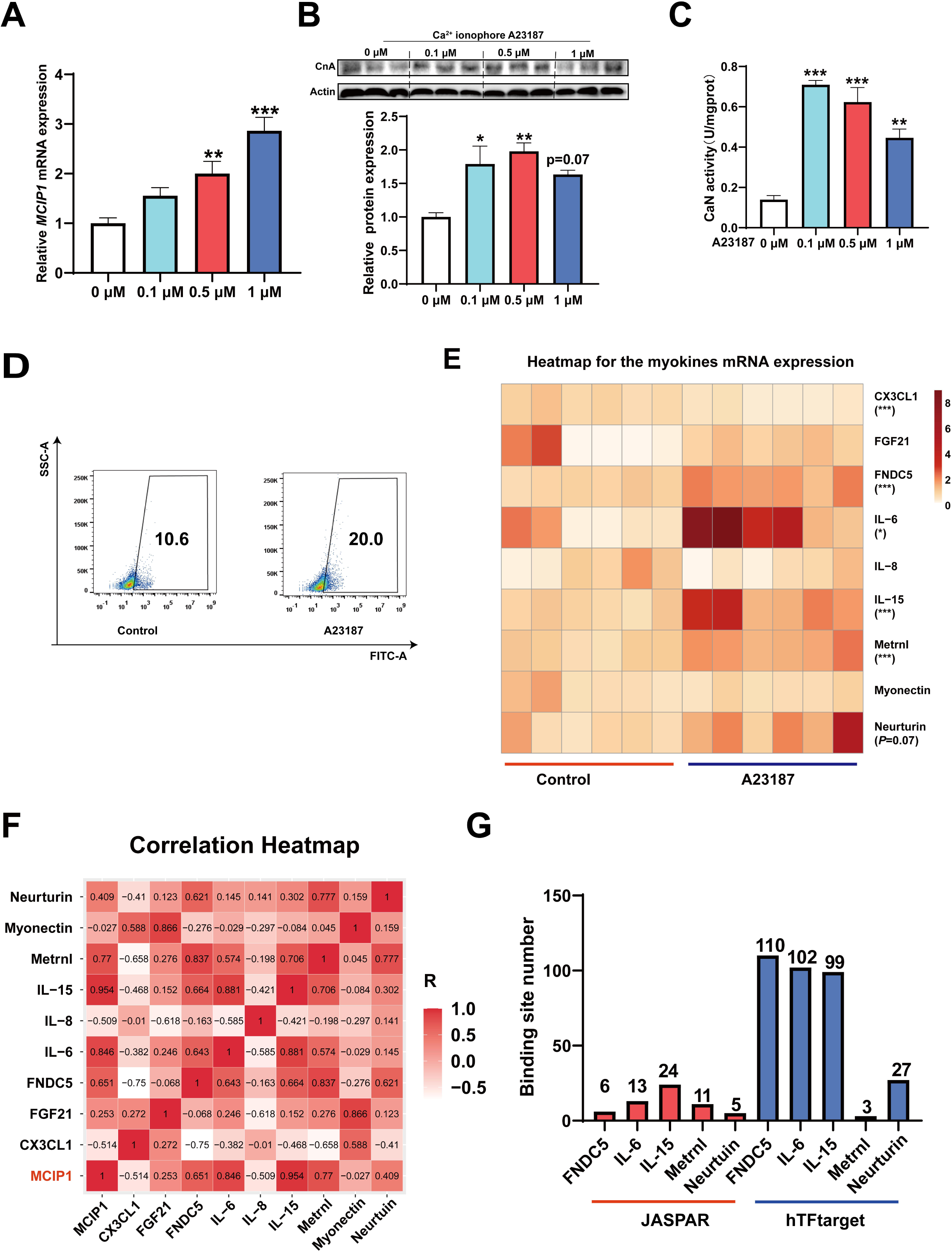
The myokines controlled by CaN. C2C12 myotubes were treated for 16 h with 0, 0.1, 0.5, and 1 μM Ca^2+^ ionophore after 2 days of differentiation. (**A)** The mRNA expression of *MCIP1*. (**B)** The protein expression of CnA. (**C)** The enzyme activity of CaN. (**D)** Fluo-4 was used to stain the Ca^2+^ and the flow cytometry assay was used to detect Ca^2+^ fluorescence in C2C12 myotubes in control and 0.5 μM A23187 groups. (**E)** The heatmap for the myokines mRNA expression in control and 0.5 μM A23187 groups. Color gradient represents relative mRNA expression with darker colors indicating higher expression. **(F)** Correlation analysis of gene expression values of myokines and MCIP1 gene performed by linear regression with Pearson’s correlation coefficient (r). Color gradient represents correlation coefficient with darker colors indicating higher positive correlation. (**G)** The number of binding sites for transcription factors NFATc1 were predicted by hTFtarget and JASPAR. For A, N=6 per group. For B-D, N=3 per group. For E, N=4 per group. \**P* < 0.05, \*\**P* < 0.01, and \*\*\**P* < 0.001.

**Table 1.**
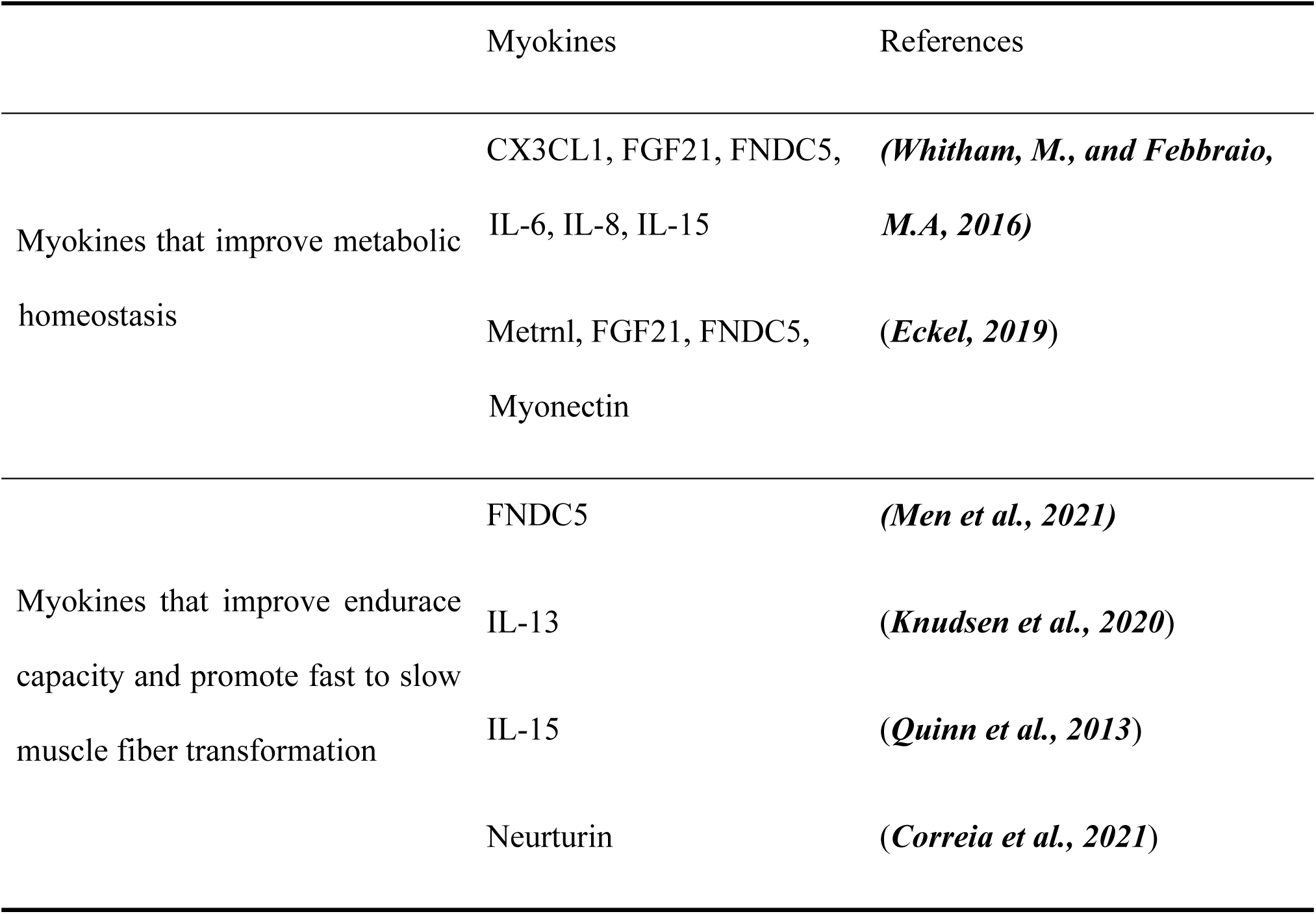
Myokines to be tested in our study.

### The myokine IL-15 expression depends on CaN/NFATc1 signaling pathway

Firstly, it was observed that treatment with 0.5 and 1 μM Ca^2+^ ionophore led to upregulation of IL-15 protein expression (Figure 7A). Subsequently, we treated C2C12 myotubes with 0.5 μM Ca^2+^ ionophore and 0.5 μM CsA to investigate their effects. We found that Ca^2+^ ionophore treatment upregulated the expression of CnA, NFATc1, and IL-15 proteins, while inhibition of CaN eliminated these effects (Figure 7B-D). Furthermore, Ca^2+^ ionophore treatment increased the expression of slow MyHC and decreased the expresssion of fast MyHC, while CsA blocked this effect (Figure 7-figure supplement 1). Based on the transcription factor motif databases (JASPAR and hIFtarget), it was predicted that a sequence (5’-AATGGAAAA-3’) in the promoter regions of IL-15 was a potential binding site of NFATc1 (Figure 7E), and DNA-protein docking analysis also revealed a high probability of binding between this sequence and the NFATc1 protein (Figure 7F). We then performed an EMSA assay to validate the binding of NFATc1 to this sequence. The probes used in the EMSA assay were shown in Figure 7G and the EMSA results were shown in Figure 7H. We observed upward migration bands when the C2C12 nuclear protein extract (NE) was incubated with the bio-NFATc1 probe (lanes 2-4), and compared to the control (lane 2) and CsA inhibition group (lane 4), Ca^2+^ treatment group showed higher expression of the band (lane 3). Moreover, the use of competing cold-NFATc1 probes showed no upward migration band (lane 5), while using the mut-NFATc1 probe again showed the band (lane 6). These findings suggest that NFATc1 may bind to the promoter of IL-15. We further investigated whether NFATc1 transcriptionally activates IL-15 through luciferase reporter assays. As shown in Figure 7I, overexpression of NFATc1 promoted NFATc1 protein expression in HEK293T cells. Additionally, after transfection with the IL-15 reporter plasmid, overexpression of NFATc1 enhanced the relative fluorescence intensity (Figure 7J), suggesting that the transcriptional activation of IL-15 is regulated by NFATc1.

**Figure 7.**
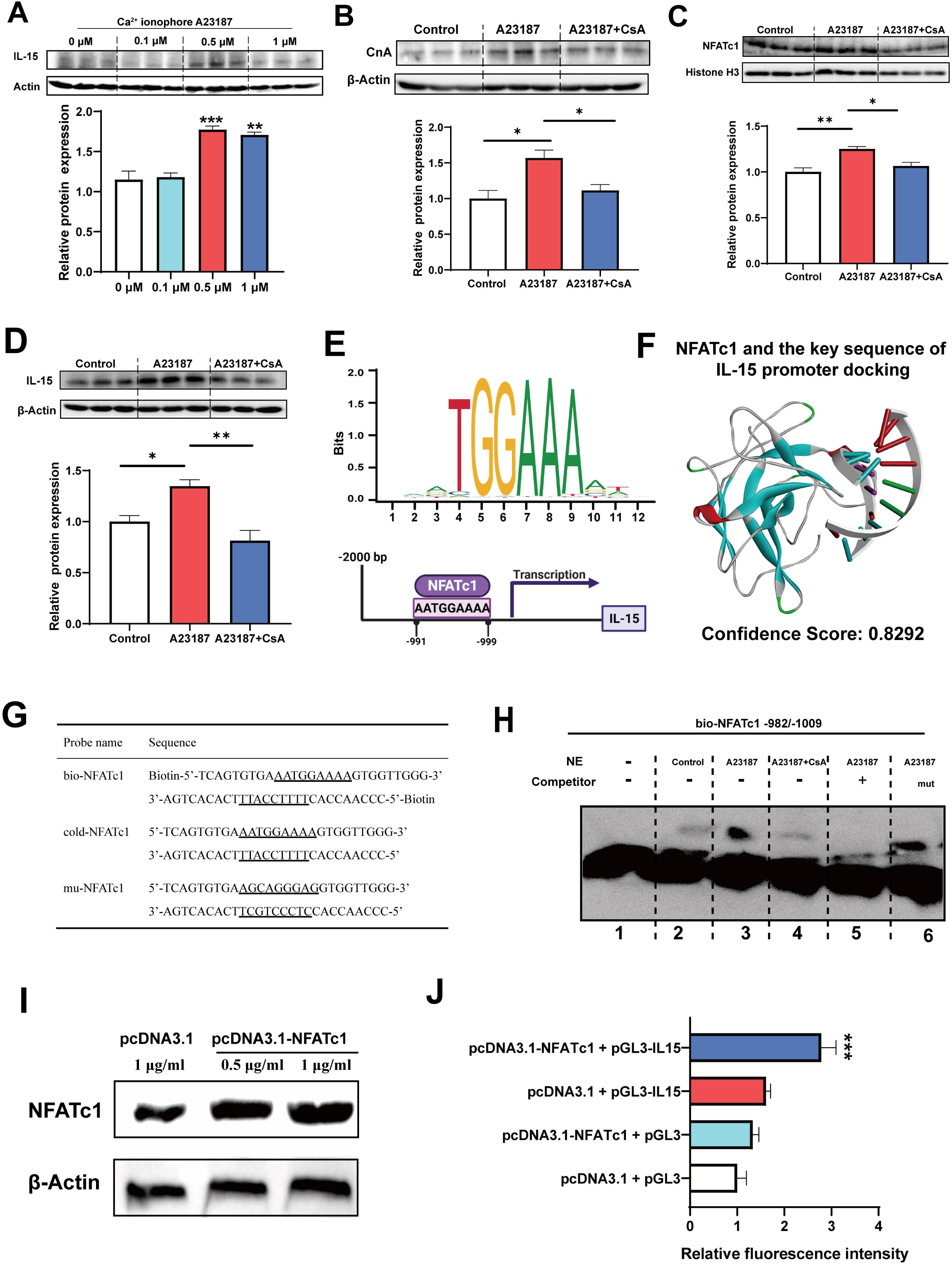
The myokine IL-15 expression depends on CaN/NFATc1 signaling pathway. (A) C2C12 myotubes were treated for 16 h with 0, 0.1, 0.5, and 1 μM Ca^2+^ ionophore after 2 days of differentiation. The protein expression of IL-15. (**B-D)** C2C12 myotubes were treated by 0.5 μM A23187 and 0.5 μM CsA for 16 h after 2 days of differentiation. The protein expression of CnA, NFATc1, and IL-15. **(E)** Sequence logo of NFATc1 motif and the predicted NFATc1 binding sites in the promoter region of IL-15. **(F)** NFATc1 and the key sequence of IL-15 (5’-AATGAAAA-3’) docking. Confidence scores above 0.7 indicate high probability of binding, scores between 0.5 and 0.7 suggest possible binding, and scores below 0.5 indicate unlikely binding. (**G)** The probe sequence of NFATc1. The underline represents the predicted binding site of NFATc1, bio-NFATc1 means oligonucleotide probes that labeled with biotin at the 5’ end, cold-NFATc1 means oligonucleotide probes that did not labeled with biotin, mu-NFATc1 means oligonucleotide probes that was mutated at the binding site. (**H)** Nuclear protein extracts (NE) with NFATc1 probe were used to EMSA assay. (**I)** The protein expression of NFATc1 after transfecting 1 μg/mL pcDNA3.1 vector, 0.5 μg/mL or 1 μg/mL pcDNA3.1-NFATc1 in HEK293T cells. (**J)** The relative luciferase intensity referred to the ratio between firefly luciferase intensity and renilla luciferase intensity. For A-D, N=3 per group. For H, N=6 per group. *P < 0.05, **P < 0.01, and ***P < 0.001.

### Eugenol promotes IL-15 level by TRPV1-mediated CaN/NFATc1 signaling pathway

We conducted further experiments to examine the effect of eugenol on IL-15 expression. Our results (Figure 8A-C) showed that EUG50 and EUG100 promoted the mRNA and protein expression of IL-15 in both GAS and TA muscle. Interestingly, in EDL and SOL muscle, which are dominated by fast and slow muscle fibers respectively, EUG100 promoted IL-15 mRNA expression (Figure 8D). We also found that the mRNA expression of IL-15 was higher in SOL muscle than in EDL muscle (Figure 8D). Moreover, Pearson’s correlation analysis showed that IL-15 expression positively correlated with MyHC I (R=0.714) and MyHC IIa (R=0.774) expression, and negatively correlated with MyHC IIb (R=-0.568) (Figure 8D). Consistent with the mRNA expression data, the IL-15 protein expression was also higher in SOL muscle than in EDL muscle (Figure 8E). Additionally, EUG50 and EUG100 increased the concentration of IL-15 in the serum (Figure 8F). These findings suggested that IL-15 was a oxidative muscle fiber type-specific myokine that was promoted by eugenol. Our in vitro experiments showed that 25 and 50 μM eugenol increased the mRNA expression and secretion of IL-15 (Figure 9A and B), which was consistent with our in vivo experiments. In addition, the inhibition of TRPV1 and CaN decreased the upregulation of eugenol on IL-15 mRNA and protein expression (Figure 9C and D). Immunofluorescence staining with IL-15 also showed similar results (Figure 9-figure supplement 1).

**Figure 8.**
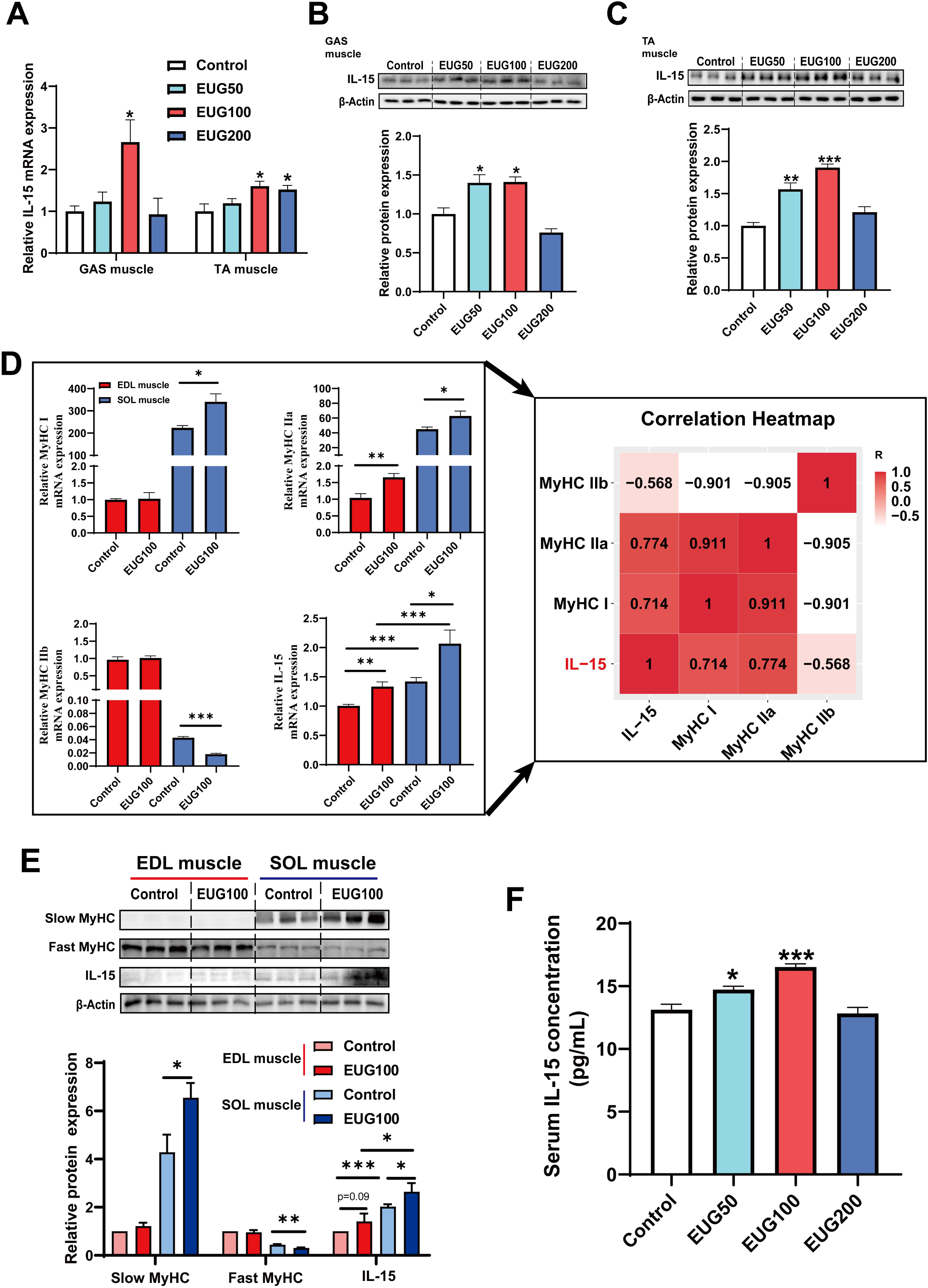
Eugenol promotes the expression and secretion of IL-15 in skeletal muscle of mice. (**A)** The *IL-15* mRNA expression in GAS and TA muscle. **(B, C)** The IL-15 protein expression in GAS and TA muscle. (**D)** Left: The mRNA expression of *MyHC I*, *MyHC IIa*, *MyHC IIb* and *IL-15* in EDL and SOL muscle of mice. Right: Correlation analysis of gene expression values performed by linear regression with Pearson’s correlation coefficient (r). Color gradient represents correlation coefficient with darker colors indicating higher positive correlation. (**E)** The protein expreesion of slow MyHC, fast MyHC, and IL-15. (**F)** The concertration of IL-15 in serum. For A, D, and F, N=6 per group. For B-C and E, N=3 per group. \**P* < 0.05, \*\**P* < 0.01, and \*\*\**P* < 0.001.

**Figure 9.**
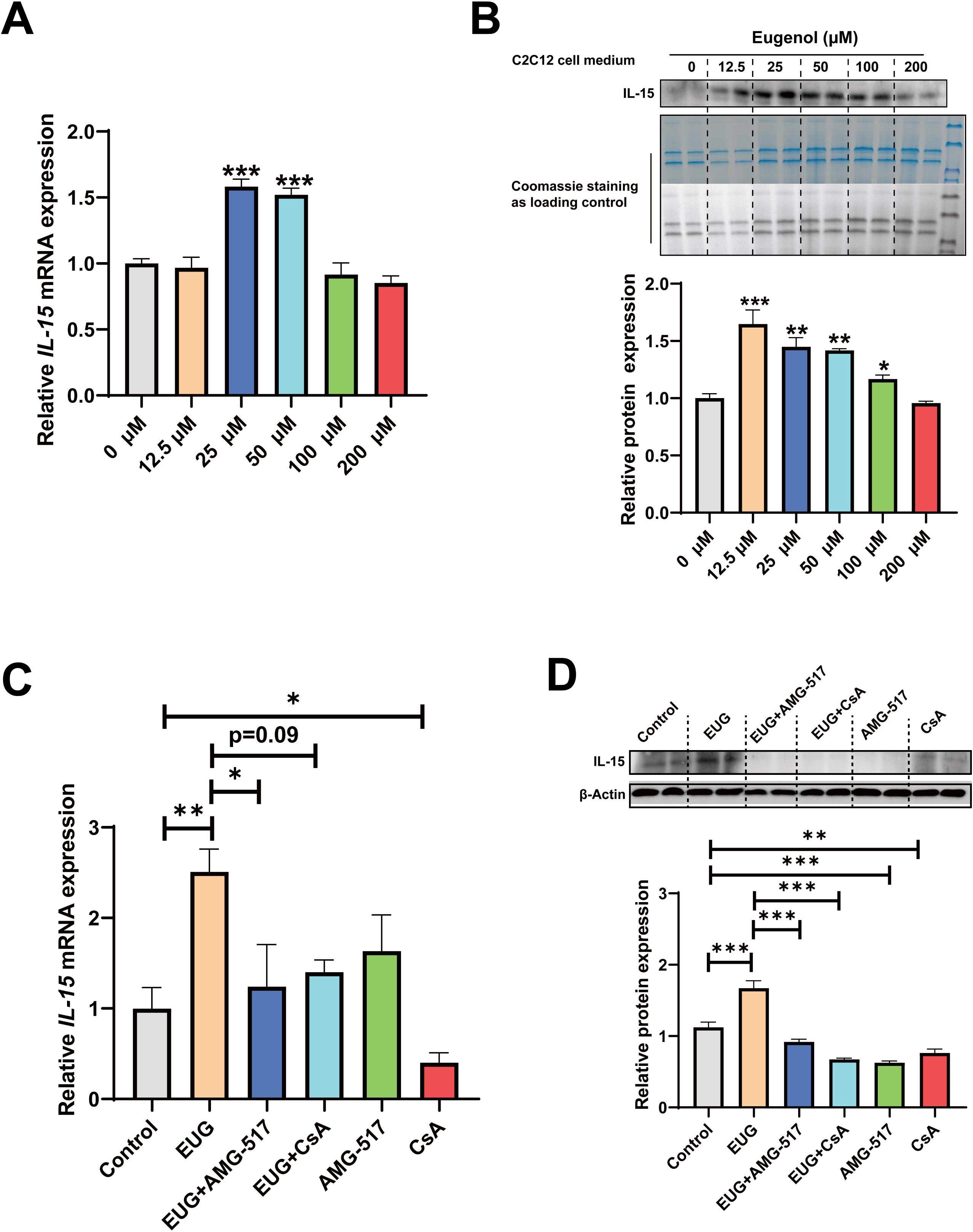
Eugenol promotes IL-15 level through TRPV1-mediated CaN/NFATc1 signaling pathway. C2C12 myotubes were treated by 0-200 eugenol for 1 day after 4 days of differentiation. (**A)** The effect of eugenol on *IL-15* mRNA expression in C2C12 myotubes. (**B)** The effect of eugenol on IL-15 protein expression in the C2C12 cell medium. Coomassie staining as loading control. **(C, D)** C2C12 myotubes were treated by 25 μM eugenol and 1 μM TRPV1 inhibitor AMG-517 or 0.5 μM CaN inhibitor CsA for 1 day after 4 days of differentiation. The mRNA and protein expression of IL-15. N=4 per group. \**P* < 0.05, \*\**P* < 0.01, and \*\*\**P* < 0.001.

## Discussion

Our study reveals that eugenol can serve as an exercise mimetic that promotes remodeling of skeletal muscle fibers and expression of the myokine IL-15 through the TRPV1-mediated CaN/NFATc1 signaling pathway. This expands the traditional biological functions of eugenol and provides a theoretical basis for its potential applications in the food and drug industry. Moreover, we identify a novel TRPV1-mediated CaN/NFATc1 signaling pathway that promotes the transformation of fast to slow muscle fibers. Importantly, we provide the first explanation of the mechanism underlying the release of the myokine IL-15.

TRPV1, like all other TRP channels, assembles as tetramers to form cation-permeable pores (***Clapham, 2003***). Initially identified as a capsaicin receptor and heat-activated ion channel that modulates pain and neurogenic inflammation (***Caterina et al., 1997***), subsequent studies have found that TRPV1 is expressed on many non-neural sites and plays roles in immunity, vasculature, obesity, and thermogenesis (***Fernandes et al., 2012***). Previous studies have also shown that capsaicin activated TRPV1 to improve endurance capacity and energy metabolism (***Luo et al., 2012***), counter obesity (***Baskaran et al., 2016***), and intervene diabetes (***Wang et al., 2012***). Therefore, TRPV1 is a strong target for the discovery of exercise mimetics, which drove us to search for TRPV1 agonists that may act as exercise mimetics. We focused on eugenol, a healthy and edible plant extract that may be a potential TRPV1 agonist. Since eugenol contains a vanilloyl fragment, it is possible to bind TRPV1 through a similar pattern to capsaicin. Indeed, previous studies have found that eugenol activates TRPV1 in a heterologous expression system and rat trigeminal ganglion neurons (***Xu et al., 2006; Yang et al., 2003***), and a recent study reported that 50 and 100 μM eugenol promoted TRPV1 expression in C2C12 myotubes (***Jiang et al., 2022***). In our study, we investigated the mRNA expression profile of TRP channels in C2C12 myotubes and skeletal muscle, and found that only TRPV1 mRNA was upregulated in response to eugenol treatment. Moreover, it was observed that the protein expression of TRPV1 was consistent with the mRNA expression. Our molecular docking results also indicated that eugenol bound to the capsaicin binding pockets of TRPV1. Based on these findings, we propose that eugenol specifically activates TRPV1 in skeletal muscle, rather than other TRP channels. This may be due to the relatively higher expression of TRPV1 in skeletal muscle compared to other TRP channels.

One of the benefits of exercise mimetics for body is to promote the skeletal muscle oxidative phenotype and endurance performance (***Fan and Evans, 2017***). Our studies showed that eugenol improved endurance performance and promoted fast-to-slow muscle fiber transformation. Slow muscle fibers are characterized by the improvement in mitochondrial function and oxidative metabolism (***Choi and Kim, 2009***). As expected, eugenol also promoted mitochondrial function and oxidative metabolism capacity in skeletal muscle. In a previous study, capsaicin-induced TRPV1 activation was shown to promote exercise endurance and the skeletal muscle oxidative phenotype (***Wang et al., 2012***). The authors suggested that TRPV1 activated calmodulin-dependent protein kinase (CaMK) by increasing intracellular Ca^2+^, and the activation of CaMK further activated PGC-1α, contributing to these effects (***Wang et al., 2012***). In skeletal muscle, the CaN signaling pathway is also an important Ca^2+^-mediated signal that promotes the muscle oxidative phenotype (***Sakuma and Yamaguchi, 2010***). CaN induces the translocation of NFATc1 to the nucleus by dephosphorylating NFATc1, thereby switching fast to slow muscle fibers (***Calabria et al., 2009***). TRPV1 activation has been shown to promote CaN activity in several other cells and tissues (***Hou et al., 2019; Ma et al., 2011; Yang et al., 2018***), but the link between TRPV1 and CaN signaling in skeletal muscle has not yet been established, nor has it been determined whether TRPV1 promotes muscle oxidative phenotype through the CaN/NFATc1 signaling pathway. Therefore, we investigated the role of TRPV1-mediated CaN signaling pathway in the regulation of muscle fiber types. We found that eugenol increased CaN expression through the activation of TRPV1, and that it promoted fast-to-slow muscle fiber transformation via the TRPV1-mediated CaN/NFATc1 signaling pathway. Recent studies have also found that eugenol increases muscle glucose uptake through the TRPV1/CaMK signaling pathway (***Jiang et al., 2022***), suggesting that both CaN and CaMK signaling may be involved in TRPV1-facilitated skeletal muscle oxidative phenotype.

Another benefit of exercise mimetics is the alleviation of obesity and improvement of metabolic health. It has been reported that an increase in slow-twitch fiber content is positively associated with reduced obesity and improved metabolic health (***Carlson et al., 2010***). Previous studies have shown that eugenol can reduce blood lipids in rats with hyperlipidemia (***Harb et al., 2019***), lower blood glucose and insulin resistance in diabetic mice (***Al-Trad et al., 2019; Sanae et al., 2014***), and reduce hepatic lipid accumulation in high-fat-fed mice (***Rodrigues et al., 2022***), demonstrating its potential in improving glucose and lipid metabolism. In our study, we found that feeding eugenol under standard diet conditions may promote fat thermogenesis and browning to facilitate fat breakdown. Interestingly, it was found that capsaicin alleviated obesity and promoted white fat browning by activating the TRPV1 (***Baskaran et al., 2016***). Our study found that eugenol promoted TRPV1 mRNA expression in adipose tissue, indicating that eugenol may promote white fat browning through TRPV1 activation.

Exercise mimetics may exert beneficial effects on the body by promoting the release of myokines (***Fan and Evans, 2017***). Some myokines, such as IL-13 (***Knudsen et al., 2020***), IL-15 (***Quinn et al., 2013***), and neurturin (***Correia et al., 2021***), have been well documented to improve metabolic homeostasis, promote fast to slow muscle fiber transformation, and improve endurance capacity. However, the regulatory mechanisms for myokines expression are largely unclear. Since myokines are usually promoted by muscle contraction, and Ca^2+^ is the main signal in muscle contraction, CaN, downstream of Ca^2+^, has been considered to regulate myokine expression. CaN has been reported to regulate the expression of myokine IL-6 (***Banzet et al., 2005; Banzet et al., 2007***). Additionally, the transcriptional activation activity of myokines IL-4 and IL-13 is thought to be promoted by transcription factor NFATc2, which is downstream of CaN (***Jacquemin et al., 2007; Zádor, 2008***). Based on the correlation and TFBS analysis, it was found that IL-15 is most likely to be regulated by the CaN/NFATc1 signaling pathway in our study. IL-15 has been reported as a myokine that improves fatty acid utilization, insulin sensitivity, and endurance capacity, and prevents obesity and diabetes (***Nadeau and Aguer, 2019***). In different cell models, CsA-mediated inhibition of CaN decreased the release of IL-15 (***Cho et al., 2002; Cho et al., 2007***). Furthermore, an increase in CaN expression in 3T3-L1 cells was accompanied by an increase in IL-15 expression (***Almendro et al., 2009***). Our study found that the CaN/NFATc1 signaling pathway contributes to the eugenol-promoted expression of IL-15. Furthermore, we observed that the expression of IL-15 was higher in the slow-twitch SOL muscle than in the fast-twitch EDL muscle, suggesting that an increase in muscle oxidative phenotype may further promote the release of IL-15. However, the mechanism underlying how IL-15 promotes the muscle oxidative phenotype remains unclear and requires further investigation.

Apparently, an interesting finding throughout our *in vitro* and *in vivo* study was that the high doses of eugenol (200 mg/kg for mice and 200 μM eugenol for C2C12 myotubes) had no effect on TRPV1-mediated CaN/NFATc1 signaling pathway, IL-15 expression, and muscle fiber type. We suspected that high doses of eugenol may cause desensitization of TRPV1 in skeletal muscle under our study conditions. This indirectly verified that the activation of TRPV1 does promote IL-15 expression and fast-to-slow muscle fiber transformation type. It is well-known that continuous activation of TRPV1 by capsaicin causes TRPV1 desensitization, which is considered to underlie the analgesic effects of capsaicin (***Koplas et al., 1997***). As previously reported, TRPV1 agonists promoted TRPV1 desensitization in a dose- and time-dependent manner by targeting lysosomes to degrade TRPV1 in HEK293 cells (***Sanz-Salvador et al., 2012***). A hypothetical model suggested that upon TRPV1 opening to allow Ca^2+^ influx, Ca^2+^/CaM binds to the CaM-binding domain of TRPV1, leading to channel inactivation, and CaN dephosphorylates TRPV1 to desensitize TRPV1 (***Hasan and Zhang, 2018***). However, most results on TRPV1 desensitization were obtained by capsaicin treatment of cells, and it is unknown whether high-dose eugenol promotes TRPV1 desensitization through a similar mechanism. Therefore, it is essential to select the appropriate dose of eugenol for the application of TRPV1 activators.

In conclusion, our findings indicate that eugenol can promote the transformation of fast to slow muscle fibers and induce the myokine IL-15 expression through the TRPV1-mediated CaN/NFATc1 signaling pathway. Moreover, our study is the first to demonstrate that the expression of IL-15 is regulated by the CaN/NFATc1 signaling pathway. These results suggest that eugenol may have potential as a novel exercise mimetic and that TRPV1 may represent a promising therapeutic target for metabolic disorders.

## Methods and Materials

### Animals, treatments, and sample collection

A total of eighty 4-week-old male C57BL/6J mice (Dashuo Experimental Animal Co.Ltd., Chengdu, China) were divided into four treatments (n = 12) using a simple randomization method. The control group were fed a basal diet supplemented with 0% eugenol (EUG, purity≥98%, Sigma, St. Louis, MO, USA), other groups were fed a basal diet supplemented with 50, 100 and 200 mg/kg eugenol, respectively (EUG50, EUG100 and EUG200). All mice were housed in individual cages (23°C ± 2°C, 12-h light/12-h dark cycle) and provided with free access to feed and water. The experiment lasted for 4 weeks. The weight of mice was measured every week. At the end of the experiment, mice anesthetized with CO_2_ were sacrificed. After being weighed and photographed, skeletal muscle [including tibialisanterior (TA), gastrocnemius (GAS), soleus (SOL), and extensor digitorum longus (EDL)], fat [including inguinal white adipose tissue (iWAT), gonadal white adipose tissue (gWAT), brown adipose tissue (BAT)] were collected and stored at -80 °C for subsequent analyses. All procedures of animal experiments were performed according to protocols approved the Animal Care Advisory Committee of Sichuan Agricultural University under permit No. YYS20200929.

### Exhausting swimming test

The forced swimming capacity test was employed in this study to evaluate the effects of eugenol on endurance capacity in mice. A total of thirty 4-week-old mice were divided into the control and EUG100 group (n=15). After four weeks of feeding, the mice with a load of lead wire (7% of body weight) attached to its tail were placed in individual swimming pools (25 ± 1°C, 35-cm depth). Exhaustive swimming time was immediately recorded when the mice failed to return to the surface continuously over a 7-second time frame and showed a lack of coordinated movements.

### Cell culture and treatments

C2C12 cells (Shanghai Cell Bank, Chinese Academy of Sciences, passages 3-8) were cultured in Dulbecco modified Eagle medium (DMEM) (Invitrogen, Carlsbad, CA, USA) supplemented with 10% fetal bovine serum (Gibco, Paisley, Scotland, UK), 100 mg/L streptomycin and 100 U/ml penicillin (Gibco) at 37°C in a 5 % CO2 atmosphere. When cells reached ∼80% confluence, 2% horse serum (Gibco) replaced 10% fetal bovine serum to induce differentiation. For eugenol treatment, cells were treated with 0, 12.5, 25, 50, 100, 200 μM EUG for 1 day after 4 days of differentiation. For Ca^2+^ ionophore A23187 (Sigma) treatment, cells were treated with 0, 0.1, 0.5, 1 μM A23187 for 16 h after 2 days of differentiation, or cells were treated with 0.5 μM A23187 and 0.5 μM CsA. For the following mechanism studies, cells were treated with 25 μM EUG and 1 μM TRPV1 inhibitor AMG-517 or 0.5 μM CaN inhibitor cyclosporin A (CsA, Sigma).

### Cell viability assay

Cell viability was analyzed using Cell-Counting Kit-8 (Beyotime, Jiangsu, China) to determine the safe dose of EUG on C2C12 myotubes. Briefly, cells were treated with EUG (0, 12.5, 25, 50, 100, 200, 400, 800, 1600 μM) for 1 day after 4 days of differentiation, 10 μL CCK-8 solution was then added to each well and then incubated for 1 h at 37 ℃. After incubation, the OD value was immediately measured at 450 nm using the SpectraMax 190 Absorbance Plate Reader.

### Measurement of intracellular calcium ion (Ca^2+^)

After 1 day of treatment with C2C12 myotubes in 24-well cell culture plates, the cells were digested using Trypsin-EDTA solution and transferred from each well into a centrifuge tube. The Trypsin-EDTA solution was removed and the cells were washed 3 times with a calcium-free PBS solution. Following this, the PBS solution was removed and 200 μL of 5 μM Ca^2+^ fluorescent probe Fluo-4 (Beyotime) was added to the cells and incubated at 37°C for 30 minutes. After the incubation, the probe was removed by washing the cells 3 times with PBS. Finally, Ca^2+^ fluorescence was detected using flow cytometry (FACSVerse, BD Biosciences, East Rutherford, NJ, USA) and analyzed using FlowJo 10.0.7 software.

### Gene expression and mitochondrial DNA quantitative PCR (qPCR)

Total RNA was extracted using RNA isolater Total RNA Extraction Reagent (Vazyme, Nanjing, China) and genomic DNA was extracted using mammalian genomic DNA extraction kit (Beyotime) according to the instructions. After measuring the concentration of total RNA, total RNA reverse transcribed to cDNA using HiScript II Q RT Supermix (Vazyme). qPCR was performed using ChamQ SYBR Color qPCR Master Mix (Vazyme) on a 7900 HT Real-time PCR system (384-cell standard block) (Applied Biosystems). Relative mtDNA was quantified by qPCR using primers for mitochondrially-encoded *Nd1* normalized to nuclear-encoded *36B4 (Rplp0)* DNA. And *GAPDH* was used as an endogenous control for normal qPCR. The primer sequences are listed in Appendix 1—table 1.

### Protein extraction and Western Blot

RIPA lysis buffer (Beyotime) was used to extract total protein. The nuclear protein was extracted using NE-PER Nuclear and Cytoplasmic Extraction Reagents (ThermoFisher). Protein from conditioned media was extracted using methanol-choloroform precipitation method as previous (***Jakobs et al., 2013***). Protein concentration was determined by the BCA assays, and then proteins were transferred to a nitrocellulose membrane using a wet Trans-Blot system (Bio-Rad, Hercules, CA, USA). The primary antibodies used were anti-slow MyHC (Sigma, cat. no. M8421), anti-fast MyHC (Sigma, cat. no. M4276), anti-TRPV1 (Alomone, cat. no. ACC030), anti-Calcineurin A (CnA, abcam, cat. no. ab90540), anti-NFATc1 (Cell Signaling Technology, cat. no. #8032), anti-IL-15 (R&D system, cat. no. AF447), anti-PGC-1α (Proteintech, cat. no. 66369-1-Ig). anti-NDUFA9 (GeneTex, cat. no. GTX132978), anti-SDHA (GeneTex, cat. no. GTX636098), anti-UQCRC1 (GeneTex, cat. no. GTX630393), anti-MTCO1(Bioss, cat. no. bs-3953R), anti-ATP5B (GeneTex, cat. no. GTX132925), anti-FABP1 (Cell Signaling Technology, cat. no. #13368), anti-UCP-1 (Proteintech, Cat. no. 23673-1-AP), anti-PRDM16 (R&D system, cat. no. AF6295), anti-β-actin (TransGen, cat. no. HC201-01), and anti-Histone H3 (Beyotime, cat. no. AF0009). Coomassie staining is depicted as loading control for conditioned media protein (***Welinder and Ekblad, 2011***). The signal was visualized using a Clarity Western ECL Substrate (BioRad, Hercules, CA, USA) and a ChemiDoc XRS Imager System (BioRad). The target band density was identified using Image Lab 5.1 (BioRad).

### Immunofluorescence (IF)

After treatment, C2C12 myotubes were washed three times (5 min each time) with phosphate-buffered saline (PBS) and fixed in Immunol Staining Fix Solution (Beyotime) for 20 min. Then C2C12 myotubes were permeabilized with 0.5% Triton X-100 for 20 min, blocked with blocking buffer for 2 hours at 37 ℃, incubated with the primary antibodies including slow MyHC (1:50, Sigma, Cat. No. M8421) for 16 hours, and incubated with the fluorescent secondary antibody (1:1000, Cell Signaling, USA) for 2 hours at 37 ℃. Finally, 4,6-diamidino-2-phenylindole (DAPI) (Beyotime) was used to stain cell nucleus for 10 minutes at room temperature. A positive signal was detected and captured using fluorescence microscopy (Lecia DMI4000 B).

### Molecular-protein docking and DNA-protein docking

The protein structures of TRPV1 (PDB codes: 5IS0) and NFATc1 (PDB codes: 1A66) were downloaded from the Research Collaboratory for Structural Bioinformatics Protein Data Bank database (RCSB PDB, https ://www.rcsb.org/) (***Burley et al., 2019***). The structure of eugenol (ZINC1411) was downloaded from the ZINC database (https://zinc.docking.org/substances/). The DNA structure was generated using Discovery Studio 2019 software (Discovery Studio 2019; BIOVIA; San Diego, USA). Discovery studio 2019 software was used to perform molecular-protein docking. HDOCK (http://hdock.phys.hust.edu.cn/) was used to perform DNA-protein docking.

### Prediction of transcription factors binding sites (TFBS)

The gene promoter sequences of mouse (upstream 2 kb) were acquired from National Center for Biotechnology Information. hIFtarget (http://bioinfo.life.hust.edu.cn/hTFtarget#!/) and JASPAR (http://jaspar.ge-nereg.net/) applied to predict the potential NFATc1 binding sites at the promoter of genes. A putative binding sites predicted by both two tools were selected for further electrophoretic mobility shift (EMSA) probes design.

### Electrophoretic mobility shift assay (EMSA)

After synthesizing the single-stranded probes for EMSA, annealing buffer for DNA oligos (5x) (Beyotime) was used to anneal to form double-stranded DNA probes. EMSA assay were performed using chemiluminescent EMSA Kit (Beyotime) according to the instructions. Briefly, 10 µL reaction system with nuclease-free water, EMSA/Gel-Shift Binding Buffer, nuclear protein, and probe was transferred to a nitrocellulose membrane using a wet Trans-Blot system. The reaction system includes negative control reaction (no nuclear protein), sample reaction, probe cold competition reaction (100-fold unlabeled probe), mutation probe cold competition reaction (100-fold unlabeled mutation probe). The signal was visualized using a Clarity Western ECL Substrate (BioRad, Hercules, CA, USA) and a ChemiDoc XRS Imager System (BioRad).

### Plasmid construction and extraction

The plasmid of pcDNA3.1 vector, pCDNA3.1-NFATc1 (The NFATc1 coding sequences were inserted into the pcDNA3.1 vector), pGL3 vector, and pGL3-IL15 (The sequence of 2 kb upstream of the IL-15 promoter sequence were inserted into the pGL3 vector) were from Tsingke Biotechnology Co., Ltd (Beijing, China). Plasmid amplification is provided in *Escherichia coli* bacteria cells. Plasmid Maxi Preparation Kit for All Purpose (Beyotime) was used to extract plasmids.

### Dual-luciferase reporter gene assay

HEK293T cells in 48-well plates were transfected with pcDNA3.1 together with pGL3 or pcDNA3.1-NFATc1 together with pGL3 or pcDNA3.1 together with pGL3-IL15 or pcDNA3.1-NFATC1 together with pGL3-IL15. Luciferase activity was detected by a Dual-Luciferase assay kit (Beyotime) with Glomax 96 microplate luminometer (Promega) in luminometer mode. The raw values of firefly luciferase were normalized to Renilla luciferase.

### Enzyme activities analysis

The tissue and C2C12 myotube were homogenized in saline and centrifuged at 3500 × g 4 ℃ for 10 minutes. The supernatant was carefully transferred to a centrifuge tube. Total protein concentration was determined by the BCA assays. The enzyme activity of lactate dehydrogenase (LDH), malate dehydrogenase (MDH), succinic dehydrogenase (SDH), and CaN were measured using commercial assay kits (Nanjing Jiancheng Bioengineering Institute, Nanjing, China).

### Statistical analyses

SAS 9.4 software was used to perform one-way ANOVA and t test. After the normality and homogeneity test, one-way ANOVA followed by Duncan’s multiple range test was performed for multiple-groups comparisons. Student’s t test was performed for two-groups comparisons. Correlation analysis was performed by Pearson’s correlation coefficient analysis. Statistical methods were not used to predetermine sample size. GraphPad Prism 8.0 (GraphPad Software, Inc., San Diego, CA) software was used to draw column charts and Omicstudio tools (https://www.omicstudio.cn/tool.) was used to draw heatmaps. Data were expressed as mean ± SEM. Statistical significance was defined as ^#^*P*< 0.1, \**P* < 0.05, \*\**P* < 0.01, and \*\*\**P* < 0.001 for all figures.

## Author contributions

Tengteng Huang: Investigation, Data curation, Formal analysis, Writing - original draft. Xiaoling Chen: Conceptualization, Methodology, Project administration, Supervision. Jun He: Methodology. Ping Zheng: Methodology. Yuheng Luo: Methodology. Aimin Wu: Methodology. Hui Yan: Methodology. Bing Yu: Methodology. Daiwen Chen: Methodology. Zhiqing Huang: Conceptualization, Funding acquisition, Methodology, Supervision, Writing - review & editing

## Data availability

Data will be made available on request.

## Source data files

**Figure 1-Source Data 1. Western blot raw image.**

**Figure 2-Source Data 1. Western blot raw image.**

**Figure 3-Source Data 1. Western blot raw image.**

**Figure 4-Source Data 1. Western blot raw image.**

**Figure 5-Source Data 1. Western blot raw image.**

**Figure 6-Source Data 1. Western blot raw image.**

**Figure 7-Source Data 1. Western blot raw image.**

**Figure 8-Source Data 1. Western blot raw image.**

**Figure 9-Source Data 1. Western blot raw image.**

---

**Figure 1-figure supplement 1.**
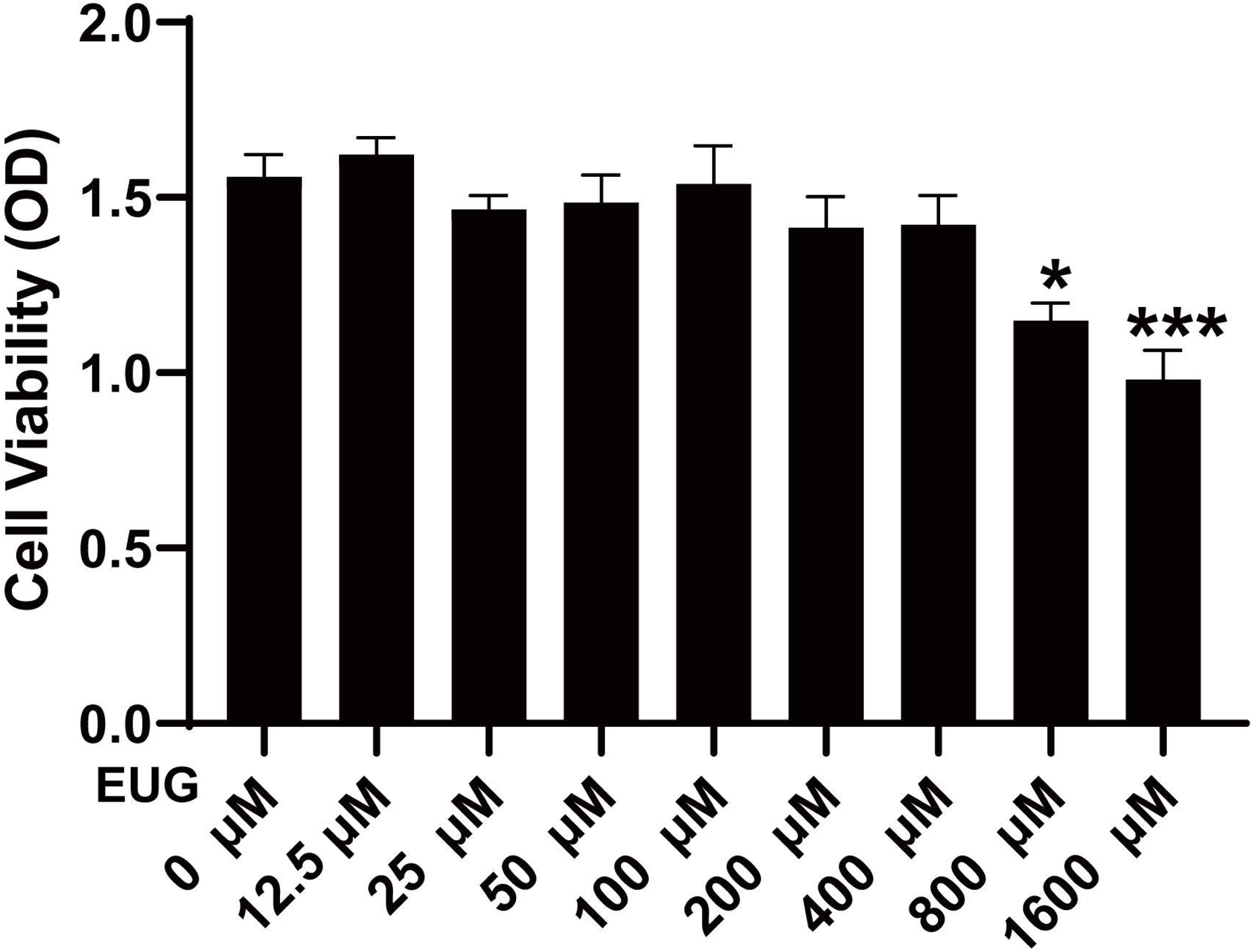
Effect of eugenol on C2C12 cell viability. Cells were treated with 0, 12.5, 25, 50, 100, 200 μM EUG for 1 day after 4 days of differentiation. The C2C12 cell viability was measured using CCK-8 kit. N=6 per group. *P < 0.05, **P < 0.01, and ***P < 0.001.

**Figure 4-figure supplement 1.**
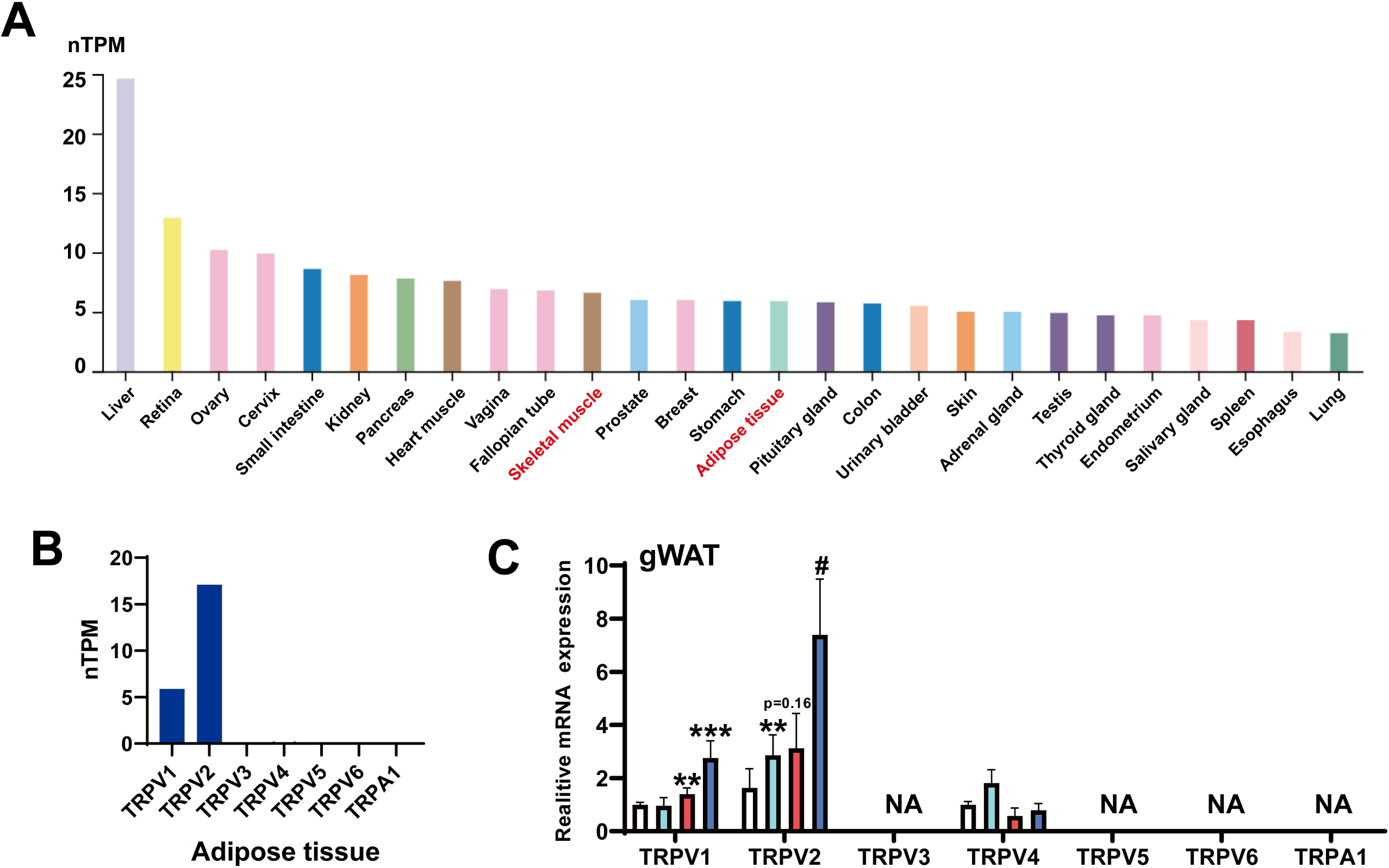
TRP channels expression profiles and *TRPV1* mRNA eexpression in adipose tissue. The gene expression profile was obtained from the GTEx dataset in The Human Protein Atlas (https://www.proteinatlas.org/). (**A)** TRPV1 expression profiles in tissues. (**B)** TRP channels expression profiles in adipose tissue. (**C)** The mRNA expression of TRP channels in adipose tissue. For C, N=6 per group. ^#^*P*<0.1, \**P* < 0.05, \*\**P* < 0.01, and \*\*\**P* < 0.001.

**Figure 4-figure supplement 2.**
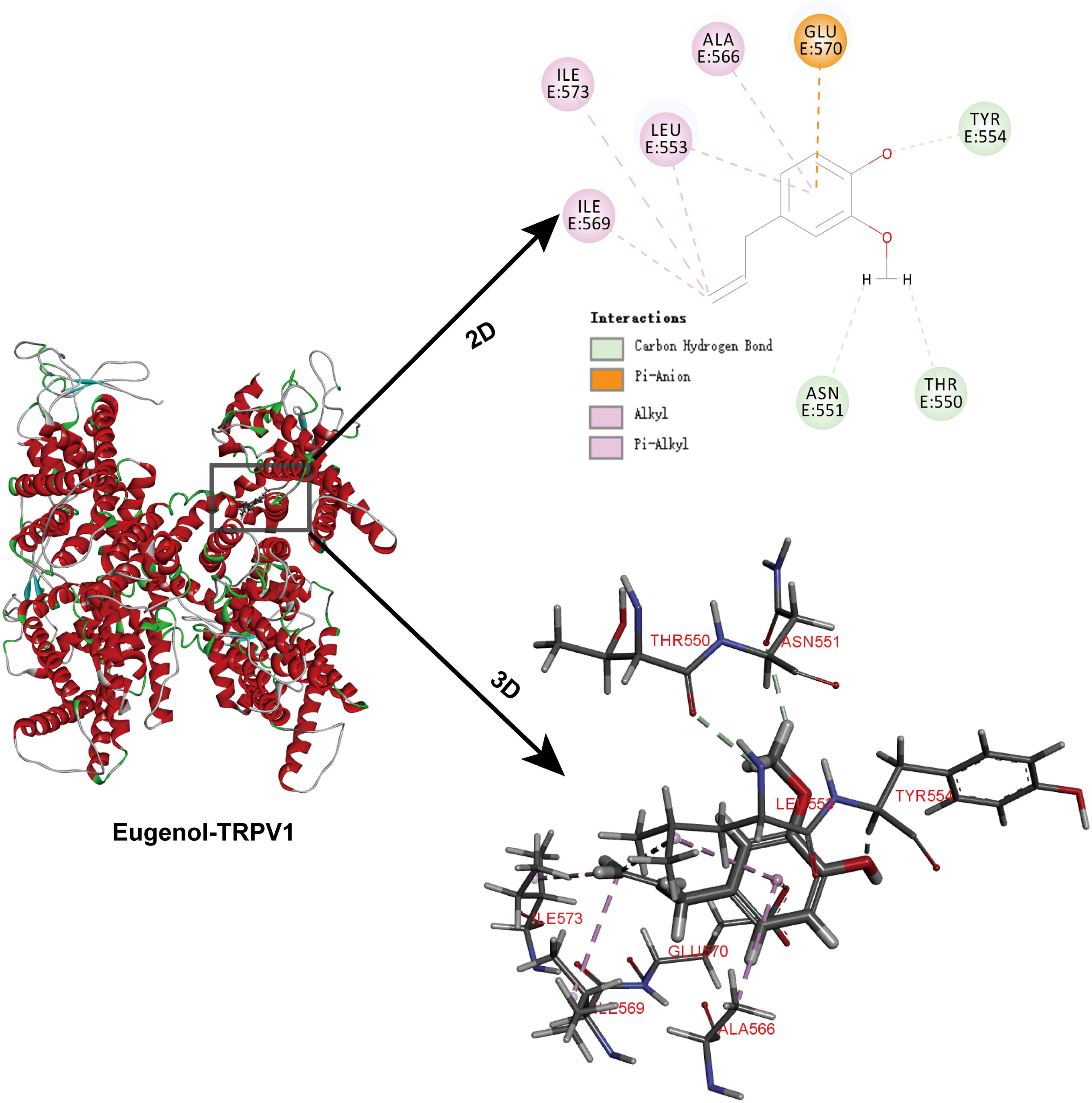
Molecular docking for eugenol and TRPV1. The capsaicin binding sites (TYR511, SER512, THR550, and GLU570) were selected as the binding pocket. The figure showed TRPV1 amino acid residues interacting with eugenol and the intermolecular force.

**Figure 4-figure supplement 3.**
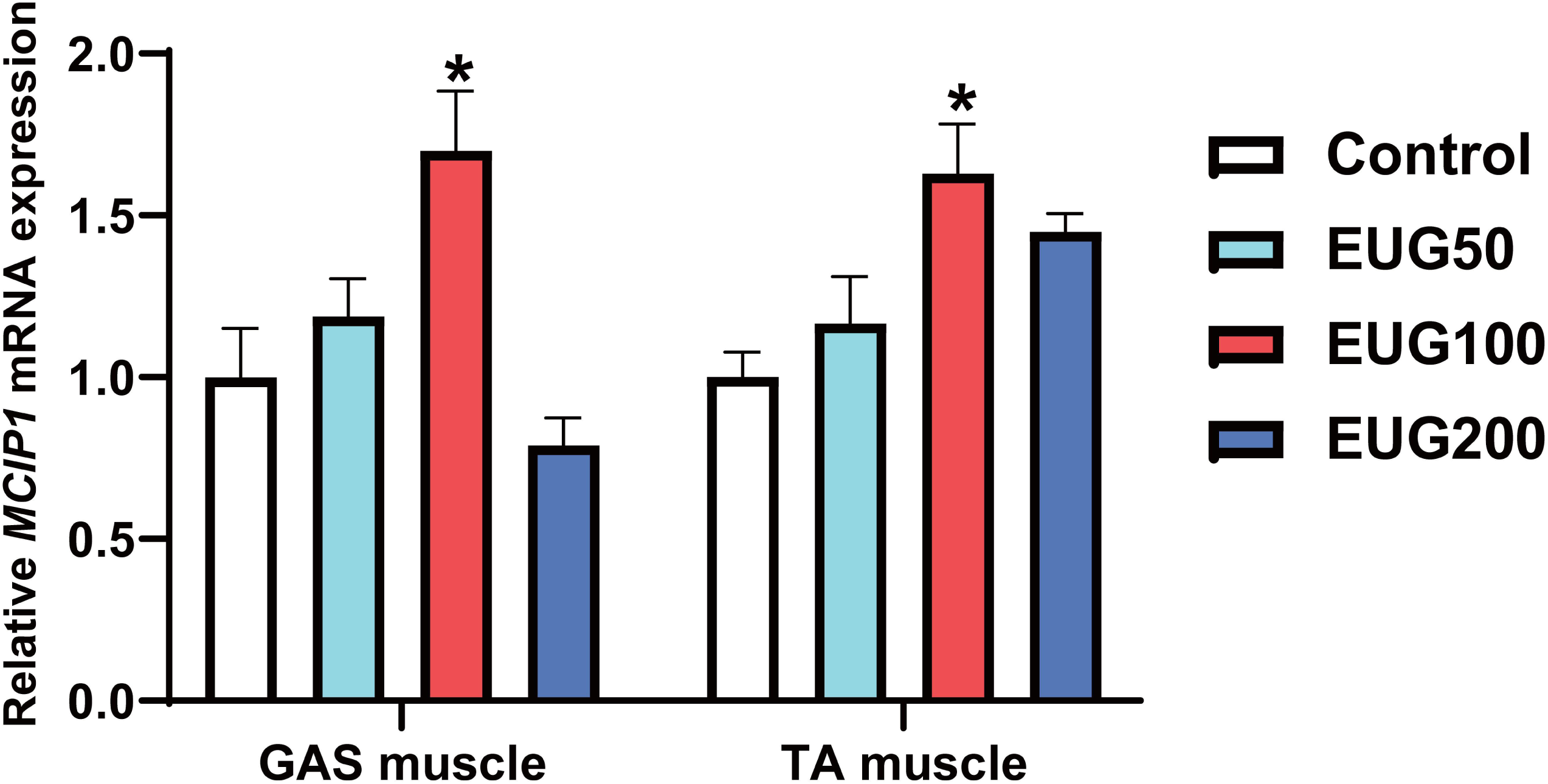
The mRNA expression of *MCIP1*. N=6 per group. *P < 0.05, **P < 0.01, and ***P < 0.001.

**Figure 7-figure supplement 1.**
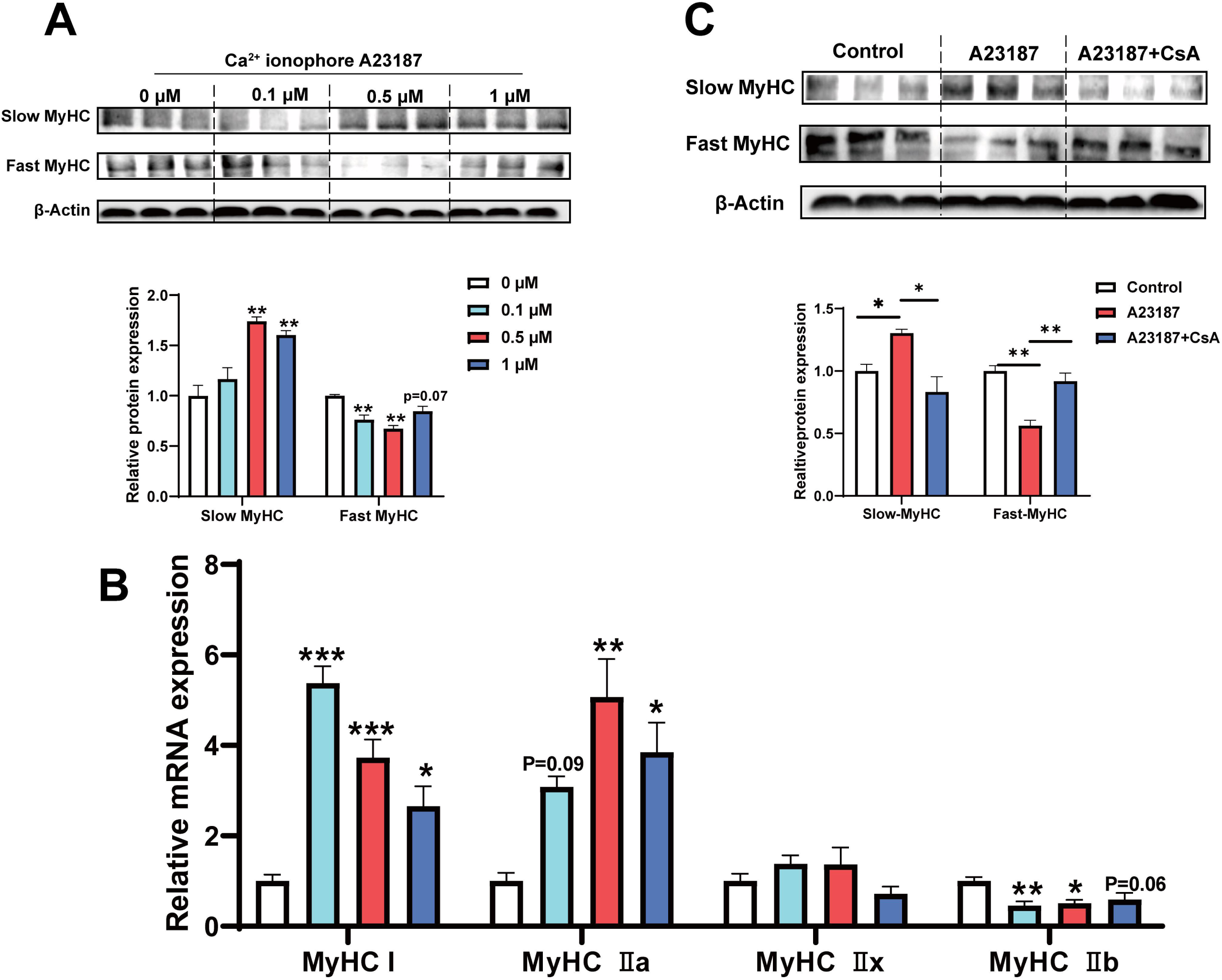
Ca^2+^ ionophore A23187 promotes the transformation of fast to slow muscle fiber by CaN signaling pathway. **A-C** The mRNA and protein expression of muscle fiber type in C2C12 myotubes. For B, N=4 per group. For A and C, N=3 per group. \**P* < 0.05, \*\**P* < 0.01, and \*\*\**P* < 0.001.

**Figure 9-figure supplement 1.**
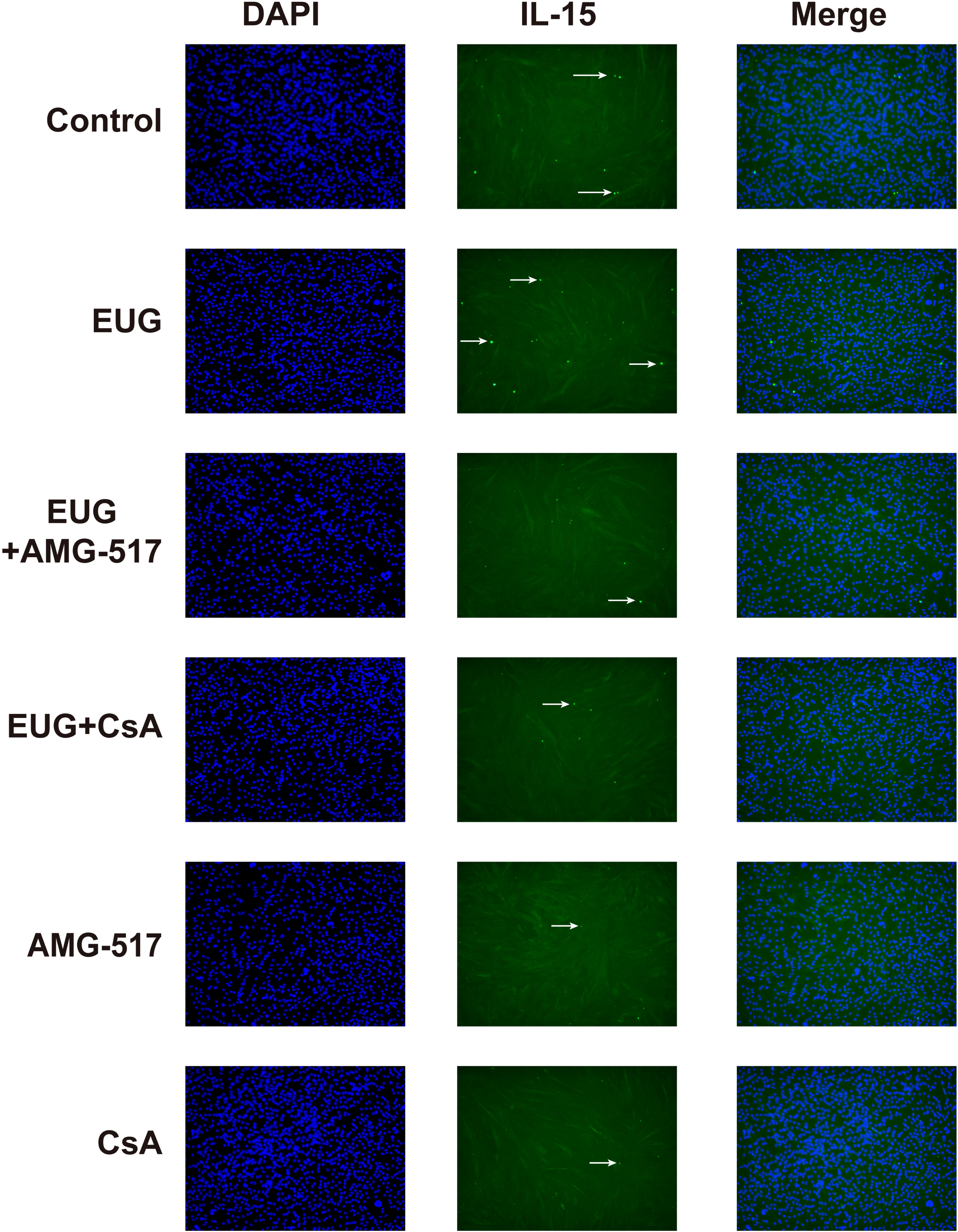
Representative immunofluorescence images of IL-15. IL-15 (Green fluorescence) and DAPI (Blue fluorescence). Magnification: × 200.

## References

1. Almendro V, Fuster G, Ametller E, Costelli P, Pilla F, Busquets S, Figueras M, Argilés JM, López-Soriano FJ. 2009. Interleukin-15 increases calcineurin expression in 3T3-L1 cells: possible involvement on in vivo adipocyte differentiation. International Journal of Molecular Medicine 24:453–458. doi: 10.3892/ijmm_00000252.

2. Al-Trad B, Alkhateeb H, Alsmadi W, Al-Zoubi M. 2019. Eugenol ameliorates insulin resistance, oxidative stress and inflammation in high fat-diet/streptozotocin-induced diabetic rat. Life Sciences 216:183–188. doi: 10.1016/j.lfs.2018.11.034.

3. Banzet S, Koulmann N, Sanchez H, Serrurier B, Peinnequin A, Alonso A, Bigard X. 2007. Contraction-induced interleukin-6 transcription in rat slow-type muscle is partly dependent on calcineurin activation. Journal of Cellular Physiology 210:596–601. doi: 10.1002/jcp.20854.

4. Banzet S, Koulmann N, Simler N, Birot O, Sanchez H, Chapot R, Peinnequin A, Bigard X. 2005. Fibre-type specificity of interleukin-6 gene transcription during muscle contraction in rat: association with calcineurin activity. The Journal of Physiology 566:839–847. doi: 10.1113/jphysiol.2005.089193.

5. Baskaran P, Krishnan V, Ren J, Thyagarajan B. 2016. Capsaicin induces browning of white adipose tissue and counters obesity by activating TRPV1 channel-dependent mechanisms. British Journal of Pharmacology 173:2369–2389. doi: 10.1111/bph.13514.

6. Burley SK, Berman HM, Bhikadiya C, Bi C, Chen L, Di Costanzo L, Christie C, Dalenberg K, Duarte JM, Dutta S, Feng Z, Ghosh S, Goodsell DS, Green RK, Guranovic V, Guzenko D, Hudson BP, Kalro T, Liang Y, Lowe R, Namkoong H, Peisach E, Periskova I, Prlic A, Randle C, Rose A, Rose P, Sala R, Sekharan M, Shao C, Tan L, Tao Y-P, Valasatava Y, Voigt M, Westbrook J, Woo J, Yang H, Young J, Zhuravleva M, Zardecki C. 2019. RCSB Protein Data Bank: biological macromolecular structures enabling research and education in fundamental biology, biomedicine, biotechnology and energy. Nucleic Acids Research 47:D464–D474. doi: 10.1093/nar/gky1004.

7. Calabria E, Ciciliot S, Moretti I, Garcia M, Picard A, Dyar KA, Pallafacchina G, Tothova J, Schiaffino S, Murgia M. 2009. NFAT isoforms control activity-dependent muscle fiber type specification. Proceedings of the National Academy of Sciences of the United States of America 106:13335–13340. doi: 10.1073/pnas.0812911106.

8. Carlson SA, Fulton JE, Schoenborn CA, Loustalot F. 2010. rend and prevalence estimates based on the 2008 physical activity guidelines for Americans. American Journal of Preventive Medicine 39:305–313. doi: 10.1016/j.amepre.2010.06.006.

9. Carnevale V, Rohacs T. 2016. TRPV1: A target for rational drug design. Pharmaceuticals 9. doi: 10.3390/ph9030052.

10. Caterina MJ, Schumacher MA, Tominaga M, Rosen TA, Levine JD, Julius D. 1997. The capsaicin receptor: a heat-activated ion channel in the pain pathway. Nature 389:816–824. doi: 10.1038/39807.

11. Cho M-L, Ju JH, Kim K-W, Moon Y-M, Lee S-Y, Min S-Y, Cho Y-G, Kim H-S, Park K-S, Yoon C-H, Lee SH, Park S-H, Kim H-Y. 2007. Cyclosporine A inhibits IL-15-induced IL-17 production in CD4+ T cells via down-regulation of PI3K/Akt and NF-kappaB. Immunology Letters 108:88–96. doi: 10.1016/j.imlet.2006.11.001.

12. Cho M-L, Kim W-U, Min S-Y, Min D-J, Min J-K, Lee S-H, Park S-H, Cho C-S, Kim H-Y. 2002. Cyclosporine differentially regulates interleukin-10, interleukin-15, and tumor necrosis factorproduction by rheumatoid synoviocytes. Arthritis & Rheumatism 46:42–51. doi: 10.1002/1529-0131(200201)46:1<42::AID-ART10026>3.0.CO;2-A.

13. Choi YM, Kim BC. 2009. Muscle fiber characteristics, myofibrillar protein isoforms, and meat quality. Livestock Science 122:105–118. doi: 10.1016/j.livsci.2008.08.015.

14. Clapham DE. 2003. TRP channels as cellular sensors. Nature 426:517–524. doi: 10.1038/nature02196.

15. Correia JC, Kelahmetoglu Y, Jannig PR, Schweingruber C, Shvaikovskaya D, Zhengye L, Cervenka I, Khan N, Stec M, Oliveira M, Nijssen J, Martínez-Redondo V, Ducommun S, Azzolini M, Lanner JT, Kleiner S, Hedlund E, Ruas JL. 2021. Muscle-secreted neurturin couples myofiber oxidative metabolism and slow motor neuron identity. Cell Metabolism 33:2215–2230.e8. doi: 10.1016/j.cmet.2021.09.003.

16. Devi KP, Nisha SA, Sakthivel R, Pandian SK. 2010. Eugenol (an essential oil of clove) acts as an antibacterial agent against Salmonella typhi by disrupting the cellular membrane. Journal of Ethnopharmacology 130:107–115. doi: 10.1016/j.jep.2010.04.025.

17. Duan Y, Li F, Tan B, Yao K, Yin Y. 2017. Metabolic control of myofibers: promising therapeutic target for obesity and type 2 diabetes. Obesity Reviews 18:647–659. doi: 10.1111/obr.12530.

18. Eckel J. 2019. Myokines in metabolic homeostasis and diabetes. Diabetologia 62:1523–1528. doi: 10.1007/s00125-019-4927-9.

19. Egan B, Zierath JR. 2013. Exercise metabolism and the molecular regulation of skeletal muscle adaptation. Cell Metabolism 17:162–184. doi: 10.1016/j.cmet.2012.12.012.

20. Fan W, Evans RM. 2017. Exercise Mimetics: Impact on Health and Performance. Cell Metabolism 25:242–247. doi: 10.1016/j.cmet.2016.10.022.

21. Feige JN, Lagouge M, Canto C, Strehle A, Houten SM, Milne JC, Lambert PD, Mataki C, Elliott PJ, Auwerx J. 2008. Specific SIRT1 activation mimics low energy levels and protects against diet-induced metabolic disorders by enhancing fat oxidation. Cell Metabolism 8:347–358. doi: 10.1016/j.cmet.2008.08.017.

22. Fernandes ES, Fernandes MA, Keeble JE. 2012. The functions of TRPA1 and TRPV1: moving away from sensory nerves. British Journal of Pharmacology 166:510–521. doi: 10.1111/j.1476-5381.2012.01851.x.

23. Harb AA, Bustanji YK, Almasri IM, Abdalla SS. 2019. Eugenol Reduces LDL Cholesterol and Hepatic Steatosis in Hypercholesterolemic Rats by Modulating TRPV1 Receptor. Scientific Reports 9:14003. doi: 10.1038/s41598-019-50352-4.

24. Hasan R, Zhang X. 2018. Ca2+ regulation of TRP ion channels. International journal of molecular Sciences 19. doi: 10.3390/ijms19041256.

25. Hoehn KL, Turner N, Swarbrick MM, Wilks D, Preston E, Phua Y, Joshi H, Furler SM, Larance M, Hegarty BD, Leslie SJ, Pickford R, Hoy AJ, Kraegen EW, James DE, Cooney GJ. 2010. Acute or chronic upregulation of mitochondrial fatty acid oxidation has no net effect on whole-body energy expenditure or adiposity. Cell Metabolism 11:70–76. doi: 10.1016/j.cmet.2009.11.008.

26. Hou N, He X, Yang Y, Fu J, Zhang W, Guo Z, Hu Y, Liang L, Xie W, Xiong H, Wang K, Pang M. 2019. TRPV1 induced apoptosis of colorectal cancer cells by activating calcineurin-NFAT2-p53 signaling pathway. BioMed Research International 2019:6712536. doi: 10.1155/2019/6712536.

27. Jacquemin V, Butler-Browne GS, Furling D, Mouly V. 2007. IL-13 mediates the recruitment of reserve cells for fusion during IGF-1-induced hypertrophy of human myotubes. Journal of Cell Science 120:670–681. doi: 10.1242/jcs.03371.

28. Jakobs C, Bartok E, Kubarenko A, Bauernfeind F, Hornung V. 2013. Immunoblotting for active caspase-1. Methods in Molecular Biology 1040:103–115. doi: 10.1007/978-1-62703-523-1_9.

29. Jiang Y, Feng C, Shi Y, Kou X, Le G. 2022. Eugenol improves high-fat diet/streptomycin-induced type 2 diabetes mellitus (T2DM) mice muscle dysfunction by alleviating inflammation and increasing muscle glucose uptake. Frontiers in Nutrition 9. doi: 10.3389/fnut.2022.1039753.

30. Jung CH, Ahn J, Jeon T-I, Kim TW, Ha TY. 2012. Syzygium aromaticum ethanol extract reduces high-fat diet-induced obesity in mice through downregulation of adipogenic and lipogenic gene expression. Experimental and Therapeutic Medicine 4:409–414. doi: 10.3892/etm.2012.609.

31. Kaur G, Athar M, Alam MS. 2010. Eugenol precludes cutaneous chemical carcinogenesis in mouse by preventing oxidative stress and inflammation and by inducing apoptosis. Molecular Carcinogenesis 49:290–301. doi: 10.1002/mc.20601.

32. Knudsen NH, Stanya KJ, Hyde AL, Chalom MM, Alexander RK, Liou Y-H, Starost KA, Gangl MR, Jacobi D, Liu S, Sopariwala DH, Fonseca-Pereira D, Li J, Hu FB, Garrett WS, Narkar VA, Ortlund EA, Kim JH, Paton CM, Cooper JA, Lee C-H. 2020. Interleukin-13 drives metabolic conditioning of muscle to endurance exercise. Science 368:eaat3987. doi: 10.1126/science.aat3987.

33. Koplas PA, Rosenberg RL, Oxford GS. 1997. The role of calcium in the desensitization of capsaicin responses in rat dorsal root ganglion neurons. The Journal of Neuroscience 17:3525–3537. doi: 10.1523/JNEUROSCI.17-10-03525.1997.

34. Lesgards J-F, Baldovini N, Vidal N, Pietri S. 2014. Anticancer activities of essential oils constituents and synergy with conventional therapies: a review. Phytotherapy Research 28:1423–1446. doi: 10.1002/ptr.5165.

35. Luo Z, Ma L, Zhao Z, He H, Yang D, Feng X, Ma S, Chen X, Zhu T, Cao T, Liu D, Nilius B, Huang Y, Yan Z, Zhu Z. 2012. TRPV1 activation improves exercise endurance and energy metabolism through PGC-1α upregulation in mice. Cell Research 22:551–564. doi: 10.1038/cr.2011.205.

36. Ma L, Zhong J, Zhao Z, Luo Z, Ma S, Sun J, He H, Zhu T, Liu D, Zhu Z, Tepel M. 2011. Activation of TRPV1 reduces vascular lipid accumulation and attenuates atherosclerosis. Cardiovascular Research 92:504–513. doi: 10.1093/cvr/cvr245.

37. Magalhães CB, Casquilho NV, Machado MN, Riva DR, Travassos LH, Leal-Cardoso JH, Fortunato RS, Faffe DS, Zin WA. 2019. The anti-inflammatory and anti-oxidative actions of eugenol improve lipopolysaccharide-induced lung injury. Respiratory Physiology & Neurobiology 259:30–36. doi: 10.1016/j.resp.2018.07.001.

38. Men X-M, Xu Z-W, Tao X, Deng B, Qi K-K. 2021. FNDC5 expression closely correlates with muscle fiber types in porcine longissimus dorsi muscle and regulates myosin heavy chains (MyHCs) mRNA expression in C2C12 cells. PeerJ 9:e11065. doi: 10.7717/peerj.11065.

39. Meng J, Lv Z, Sun C, Qiao X, Chen C. 2020. An extract of Lycium barbarum mimics exercise to improve muscle endurance through increasing type IIa oxidative muscle fibers by activating ERRγ. FASEB Journal 34:11460–11473. doi: 10.1096/fj.202000136R.

40. Mnafgui K, Kaanich F, Derbali A, Hamden K, Derbali F, Slama S, Allouche N, Elfeki A. 2013. Inhibition of key enzymes related to diabetes and hypertension by Eugenol in vitro and in alloxan-induced diabetic rats. Archives of Physiology and Biochemistry 119:225–233. doi: 10.3109/13813455.2013.822521.

41. Nadeau L, Aguer C. 2019. Interleukin-15 as a myokine: mechanistic insight into its effect on skeletal muscle metabolism. Applied Physiology, Nutrition, and Metabolism 44:229–238. doi: 10.1139/apnm-2018-0022.

42. Narkar VA, Downes M, Yu RT, Embler E, Wang Y-X, Banayo E, Mihaylova MM, Nelson MC, Zou Y, Juguilon H, Kang H, Shaw RJ, Evans RM. 2008. AMPK and PPARδ agonists are exercise mimetics. Cell 134:405–415. doi: 10.1016/j.cell.2008.06.051.

43. Nilius B, Owsianik G, Voets T, Peters JA. 2007. Transient receptor potential cation channels in disease. Physiological Reviews 87:165–217. doi: 10.1152/physrev.00021.2006.

44. Nogueira L, Ramirez-Sanchez I, Perkins GA, Murphy A, Taub PR, Ceballos G, Villarreal FJ, Hogan MC, Malek MH. 2011. (-)-Epicatechin enhances fatigue resistance and oxidative capacity in mouse muscle. The Journal of Physiology 589:4615–4631. doi: 10.1113/jphysiol.2011.209924.

45. Pedersen BK, Febbraio MA. 2012. Muscles, exercise and obesity: skeletal muscle as a secretory organ. Nature reviews. Endocrinology 8:457–465. doi: 10.1038/nrendo.2012.49.

46. Quinn LS, Anderson BG, Conner JD, Wolden-Hanson T. 2013. IL-15 overexpression promotes endurance, oxidative energy metabolism, and muscle PPARδ, SIRT1, PGC-1α, and PGC-1β expression in male mice. Endocrinology 154:232–245. doi: 10.1210/en.2012-1773.

47. Rangwala SM, Wang X, Calvo JA, Lindsley L, Zhang Y, Deyneko G, Beaulieu V, Gao J, Turner G, Markovits J. 2010. Estrogen-related receptor gamma is a key regulator of muscle mitochondrial activity and oxidative capacity. The Journal of Biological Chemistry 285:22619–22629. doi: 10.1074/jbc.M110.125401.

48. Rodrigues M, Bertoncini-Silva C, Joaquim AG, Machado CD, Ramalho LNZ, Carlos D, Fassini PG, Suen VMM. 2022. Beneficial effects of eugenol supplementation on gut microbiota and hepatic steatosis in high-fat-fed mice. Food & Function 13:3381–3390. doi: 10.1039/d1fo03619j.

49. Sakuma K, Yamaguchi A. 2010. The functional role of calcineurin in hypertrophy, regeneration, and disorders of skeletal muscle. Journal of Biomedicine & Biotechnology 2010:721219. doi: 10.1155/2010/721219.

50. Sanae F, Kamiyama O, Ikeda-Obatake K, Higashi Y, Asano N, Adachi I, Kato A. 2014. Effects of eugenol-reduced clove extract on glycogen phosphorylase b and the development of diabetes in db/db mice. Food & Function 5:214–219. doi: 10.1039/c3fo60514k.

51. Sanz-Salvador L, Andrés-Borderia A, Ferrer-Montiel A, Planells-Cases R. 2012. Agonist- and Ca2+-dependent desensitization of TRPV1 channel targets the receptor to lysosomes for degradation. The Journal of Biological Chemistry 287:19462–19471. doi: 10.1074/jbc.M111.289751.

52. Schiaffino S, Reggiani C. 2011. Fiber types in mammalian skeletal muscles. Physiological Reviews 91:1447–1531. doi: 10.1152/physrev.00031.2010.

53. Severinsen MCK, Pedersen BK. 2020. Muscle-organ crosstalk: The emerging roles of myokines. Endocrine Reviews 41. doi: 10.1210/endrev/bnaa016.

54. Srinivasan S, Sathish G, Jayanthi M, Muthukumaran J, Muruganathan U, Ramachandran V. 2014. Ameliorating effect of eugenol on hyperglycemia by attenuating the key enzymes of glucose metabolism in streptozotocin-induced diabetic rats. Molecular and Cellular Biochemistry 385:159–168. doi: 10.1007/s11010-013-1824-2.

55. Wang P, Yan Z, Zhong J, Chen J, Ni Y, Li L, Ma L, Zhao Z, Liu D, Zhu Z. 2012. Transient receptor potential vanilloid 1 activation enhances gut glucagon-like peptide-1 secretion and improves glucose homeostasis. Diabetes 61:2155–2165. doi: 10.2337/db11-1503.

56. Welinder C, Ekblad L. 2011. Coomassie staining as loading control in Western blot analysis. Journal of Proteome Research 10:1416–1419. doi: 10.1021/pr1011476.

57. Wen W, Chen X, Huang Z, Chen D, Chen H, Luo Y, He J, Zheng P, Yu J, Yu B. 2020. Resveratrol regulates muscle fiber type conversion via miR-22-3p and AMPK/SIRT1/PGC-1α pathway. The Journal of Nutritional Biochemistry 77:108297. doi: 10.1016/j.jnutbio.2019.108297.

58. Whitham M, Febbraio MA. 2016. The ever-expanding myokinome: discovery challenges and therapeutic implications. Nature Reviews. Drug Discovery 15:719–729. doi: 10.1038/nrd.2016.153.

59. Xu H, Delling M, Jun JC, Clapham DE. 2006. Oregano, thyme and clove-derived flavors and skin sensitizers activate specific TRP channels. Nature Neuroscience 9:628–635. doi: 10.1038/nn1692.

60. Yang BH, Piao ZG, Kim Y-B, Lee C-H, Lee JK, Park K, Kim JS, Oh SB. 2003. Activation of vanilloid receptor 1 (VR1) by eugenol. Journal of Dental Research 82:781–785. doi: 10.1177/154405910308201004.

61. Yang J, Rothermel B, Vega RB, Frey N, McKinsey TA, Olson EN, Bassel-Duby R, Williams RS. 2000. Independent signals control expression of the calcineurin inhibitory proteins MCIP1 and MCIP2 in striated muscles. Circulation Research 87:E61–E68. doi: 10.1161/01.res.87.12.e61.

62. Yang Y, Guo W, Ma J, Xu P, Zhang W, Guo S, Liu L, Ma J, Shi Q, Jian Z, Liu L, Wang G, Gao T, Han Z, Li C. 2018. Downregulated TRPV1 expression contributes to melanoma growth via the calcineurin-ATF3-p53 pathway. The Journal of Investigative Dermatology 138:2205– 2215. doi: 10.1016/j.jid.2018.03.1510.

63. Zádor E. 2008. dnRas stimulates autocrine-paracrine growth of regenerating muscle via calcineurin-NFAT-IL-4 pathway. Biochemical and Biophysical Research Communications 375:265–270. doi: 10.1016/j.bbrc.2008.08.024.

