## Appendix 1-table 1 for "Eugenol mimics exercise to promote skeletal muscle fiber remodeling and myokine IL-15 expression by activating TRPV1 channel"

**Appendix 1—table 1. Primers used for real-time quantitative PCR.**

| Gene name |  | Primer Sequence (5'-3') | GeneBank ID | Product size |
| --- | --- | --- | --- | --- |
| 36B4 | Forward | ACAGATTTGTCAAGCGCATC | NM_007475.5 | 76 |
|  | Reverse | TTCCACAGGCCGATAGTAAGCA |  |  |
| ACC | Forward | ACCTGTGTGGTGGAATTTCAGT | NM_133360.3 | 79 |
|  | Reverse | ACATTCTGTTTAGCGTGGGGA |  |  |
| ATP5A1 | Forward | CATTGGTGATGGTATTGCGC | NM_007505.2 | 134 |
|  | Reverse | TCCCAAACACGACAACTCC |  |  |
| ATP5B | Forward | GAGGGCAATGATTTATACCAT | NM_016774.3 | 92 |
|  | Reverse | ATCTGTCCATATACCAACGCTA |  |  |
| ATP5G1 | Forward | GGAGTGGGAGTGCAGATTGAA | NM_001161419.1 | 136 |
|  | Reverse | TTAGATGGGGCCTCTGGTCT |  |  |
| CD36 | Forward | TGTGATCGGAACTGTGGGC | XM_030254088.1 | 80 |
|  | Reverse | ACTGGCATGAGAATGCCTCC |  |  |
| CD137 | Forward | AGTTTTGCTCCTCTACCCACA | NM_001077509.1 | 177 |
|  | Reverse | GACAGACGCCAGTACCGTTC |  |  |
| Cidea | Forward | AGCAACCAAAGAAATCGGGAA | NM_007702.2 | 108 |
|  | Reverse | ACATCTCGTACATCGTGGCTT |  |  |
| COX5B | Forward | ACCCTAATCTAGTCCCGTCC | NM_009942.2 | 89 |
|  | Reverse | CAGCCAAAACCAGATGACAG |  |  |
| COX6A1 | Forward | CATCAGGACCAAGCCCTTC | NM_007748.5 | 95 |
|  | Reverse | ACTCATCTTCATAGCCGGTCG |  |  |
| COX7A | Forward | AAGCCACTTAGAAAACCGTGT | NM_001412271.1 | 163 |
|  | Reverse | CCAGCCCAAGCAGTATAAGCA |  |  |
| CX3CL1 | Forward | CCTGACAAAGCCTGAATCCG | NM_009142.3 | 192 |
|  | Reverse | GCCTCAAAACTTCCAATGCTCT |  |  |
| Dio2 | Forward | GTGAAGATGCTCCCAATTCCA | NM_010050.4 | 69 |
|  | Reverse | CCGAGGCATAATTGTTACCTG |  |  |
| FABP4 | Forward | TTCCTTCAAACTGGGCGTGG | NM_001409513.1 | 125 |
|  | Reverse | TTGTGGTCGACTTTCCATCCC |  |  |
| FASN | Forward | GAAGAGCCTGGAAGATCGGG | XM_030245556.1 | 108 |
|  | Reverse | TTGTGGTAGAAGGACACGGC |  |  |
| FGF21 | Forward | CCTCTACACAGATGACGACCA | NM_020013.4 | 160 |
|  | Reverse | AAACCTAGAGGCTTTGACACC |  |  |
| FNDC5 | Forward | ATGAAGGAGATGGGGAGGAA | NM_027402.4 | 102 |
|  | Reverse | GCGGCAGAAGAGAGCTATAACA |  |  |
| GAPDH | Forward | AGGGCATCTTGGGCTACAC | NM_008084 | 211 |
|  | Reverse | TGGTCCAGGGTTTCTTACTCC |  |  |
| HSL | Forward | GGAGCACTACAAACGCAACG | XM_030242180.1 | 108 |
|  | Reverse | CGTTCAAATTCAGCCCCACG |  |  |
| IL-6 | Forward | CCTCTCTCTGCAAGAGACTTCCAT | NM_031168.2 | 81 |
|  | Reverse | TTGTGAAGTAGGGAAGGCCG |  |  |
| IL-8 | Forward | TGGCCCAATTACTAACAGGT | NM_011339.2 | 225 |
|  | Reverse | ACTTCACTGGAGTCCCGTA |  |  |
| IL-13 | Forward | TGGCTCTTGCTTGCCTTGGTGG | NM_008355.3 | 146 |
|  | Reverse | CCATACCATGCTGCCGTTGCA |  |  |
| IL-15 | Forward | TAGCCAGCTCATCTTCAACA | NM_001254747.1 | 104 |
|  | Reverse | GAAACACAAGTAGCACGAGA |  |  |
| MCIP1 | Forward | CCGTTGGCTGGAAACAAG | NM_019466 | 153 |
|  | Reverse | GGTCACTCTCACACACGTGG |  |  |
| Metrnl | Forward | CACCCCAACAGGACATCAGC | NM_144797.3 | 172 |
|  | Reverse | TCCTCAATGAAGCCTCGGACA |  |  |
| Myonectin | Forward | CGCATTCCACTGTCGCTTG | NM_173395.2 | 96 |
|  | Reverse | AAGGCTCCCTCAACTTCGG |  |  |
| MyHC I | Forward | CTTCTACAGGCCTGGGCTTAC | NM_080728 | 128 |
|  | Reverse | CTCCTTCTCAGACTTCCGCAG |  |  |
| MyHC IIa | Forward | TTCCAGAAGCCTAAGGTGGTC | NM_001039545 | 94 |
|  | Reverse | GCCAGCCAGTGATGTTGTAAT |  |  |
| MyHC IIx | Forward | CAACCCATACGACTACGCCT | NM_030679 | 119 |
|  | Reverse | CATCAGAAGTGAAGCCCAGAAT |  |  |
| MyHC IIb | Forward | CTTGTCTGACTCAAGCCTGCC | NM_010855 | 158 |
|  | Reverse | TCGCTCCTTTTCAGACTTCCG |  |  |
| mtND1 | Forward | AATCGCCATAGCCTTCCTAA | NC_005089.1 | 114 |
|  | Reverse | GCGTCTGCAAATGGTTGTAA |  |  |
| NDUFA1 | Forward | GTCAATCGCTACTATGTGTCC | NM_019443.2 | 117 |
|  | Reverse | TGCATAGCCTTCTAACAGGA |  |  |
| NDUFB3 | Forward | GCAACATCACCTTCCCGAGT | NM_025597.3 | 70 |
|  | Reverse | AAAGCTACCACAAACGCAGCA |  |  |
| NDUFS8 | Forward | GTTCATAGGGTCAGAGGTCAAG | NM_001271444.1 | 112 |
|  | Reverse | TCCATTAAGATGTCCTGTGCG |  |  |
| Neurturin | Forward | GGGCTACACGTCGGATGAG | NM_008738.3 | 80 |
|  | Reverse | CCAGGTCGTAGATGCGGATG |  |  |
| NRF1 | Forward | TGCTTCAGAACTGCCAACCA | NM_001410232.1 | 105 |
|  | Reverse | ATTTCACCGCCCTGTAACGT |  |  |
| NRF2 | Forward | AAAGCACAGCCAGCACATTC | NM_010902 | 86 |
|  | Reverse | TGGGATTCACGCATAGGAGC |  |  |
| PGC-1α | Forward | CCAGTACAACAATGAGCCTGC | NM_001271444.1 | 118 |
|  | Reverse | CAATCCGTCTTCATCCACG |  |  |
| POLRMT | Forward | TTGACCAGCAGAAGCAAGCC | NM_001407795.1 | 135 |
|  | Reverse | GCTGCTTTTCCTCTGAGTTCGT |  |  |
| PPARγ | Forward | TGCGATCAAAGTAGAACCTG | NM 001127330.2 | 231 |
|  | Reverse | CGGCAGTTAAGATCACACC |  |  |
| PRDM16 | Forward | CCCTGACTGTGGCAAGACCTT | [NM_027504.3](https://www.ncbi.nlm.nih.gov/entrez/viewer.fcgi?db=nucleotide&id=124107622) | 101 |
|  | Reverse | ACTTGTGGCAGACCTCGCAT |  |  |
| SDHA | Forward | CCACTCACTCTTACACACGTT | NM_023281.1 | 89 |
|  | Reverse | CCATCAGAAGATCCAGTGCAA |  |  |
| SDHB | Forward | ACCCCTTCTCTGTCTACCG | NM_001355515.1 | 130 |
|  | Reverse | AATGCTCGCTTCTCCTTGTAG |  |  |
| SDHC | Forward | TCTTCCCGCTCATGTACCAC | NM_025321.3 | 52 |
|  | Reverse | TCCCATAGCAAGTGTCGGAT |  |  |
| TBX1 | Forward | GAAACTGACCAATAACCTGCT | NM_011532.2 | 114 |
|  | Reverse | TTCTCACTGTCTTTTCGAGGG |  |  |
| TFAM | Forward | ATTTCACCGCCCTGTAACGT | NM_009360 | 193 |
|  | Reverse | TCGTTTCACACTTCGACGGAT |  |  |
| TFB1M | Forward | GAAGCACAACGCCTCGAAAG | NM_146074 | 91 |
|  | Reverse | AATGCATGAGCGAGAGGTGG |  |  |
| TFB2M | Forward | CCAATAATACGCCATTTACGTT | NM_001331055.1 | 178 |
|  | Reverse | GGCAGTAGTCATAGATCCAC |  |  |
| TMEM26 | Forward | GCACCATCACTAGAGACCAAC | NM_177794.3 | 53 |
|  | Reverse | CGTCCCCACAAACATCAGA |  |  |
| TRPA1 | Forward | TGGTTATGGAAATACCCCACT | NM_177781.5 | 111 |
|  | Reverse | ATCATGTTTCTATTTCGGAGG |  |  |
| TRPV1 | Forward | ACAGATTTGTCAAGCGCATC | NM_001001445.2 | 95 |
|  | Reverse | TTCCACAGGCCGATAGTAAGCA |  |  |
| TRPV2 | Forward | TTTTAGAGCCACTGAACAAGC | NM_011706.2 | 88 |
|  | Reverse | TAGACCAAGTAACAGGCGAA |  |  |
| TRPV3 | Forward | TCTGTGCTGGAAATCATCGTCT | NM_145099.3 | 59 |
|  | Reverse | GTCAGCATCTCATGTCGGTT |  |  |
| TRPV4 | Forward | CAACCAGCCGCACATCGTC | NM_022017.3 | 69 |
|  | Reverse | TCCTGTCGCCTCATGTCAGC |  |  |
| TRPV5 | Forward | AGTTCCGAGATGCCAACCGTA | NM_001007572.2 | 119 |
|  | Reverse | TGTCACCAACTCCCCGACCA |  |  |
| TRPV6 | Forward | AAGCCCAGGACCAATAACCG | NM_022413.4 | 195 |
|  | Reverse | TCCAAAGAAGCGAGTGACCC |  |  |
| UCP-1 | Forward | AAACACCTGCCTCTCTCGGAA | NM_009463.3 | 70 |
|  | Reverse | CCAATGAACACTGCCACACCT |  |  |
| UQCRB | Forward | TGCCTCATAGTCAGGTCCA | NM_026219.2 | 126 |
|  | Reverse | GTTAATGCGAGATGATACACT |  |  |
| UQCRC1 | Forward | ATCAAGGCACTGTCCAAGG | NM_025407.2 | 131 |
|  | Reverse | TCATTTTCCTGCATCTCCCG |  |  |
| UQCRQ | Forward | CTGATCTACACATGGGGCAAC | NM_025352.3 | 56 |
|  | Reverse | GCTGGATTCTTCCTTTTCGACT |  |  |
