## Supplementary material for "Eugenol mimics exercise to promote skeletal muscle fiber remodeling and myokine IL-15 expression by activating TRPV1 channel": Figure 1-Source Data 1

**Fig. 1D**

Fig. 1D: Slow MyHC

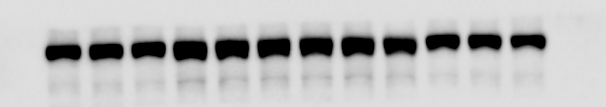

Fig. 1D: Fast MyHC

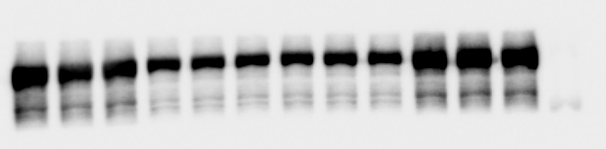

Fig. 1D:β-Actin

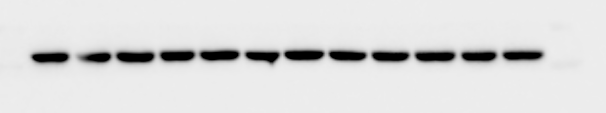

**Fig. 1E**

Fig. 1E: Slow MyHC

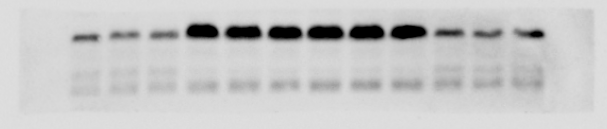

Fig. 1E: Fast MyHC

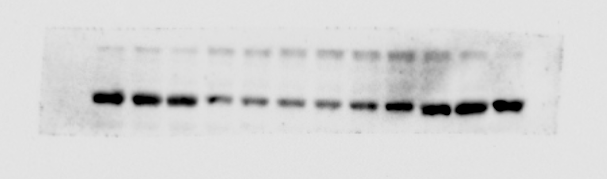

Fig. 1E:β-Actin

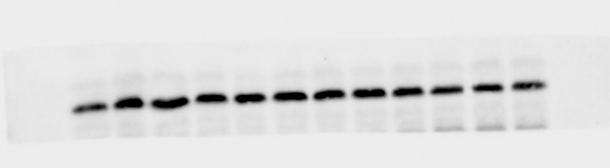

**Fig. 1F**

Fig. 1F: Slow MyHC

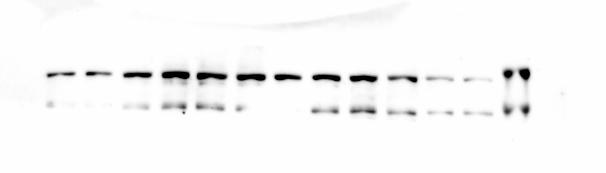

Fig. 1F: Fast MyHC

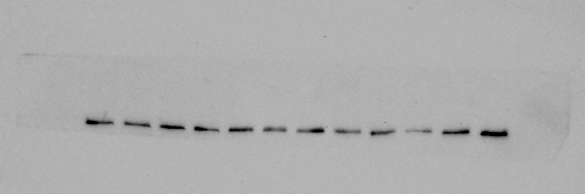

Fig. 1F:β-Actin

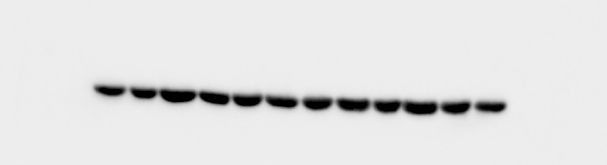
