## Supplementary material for "Eugenol mimics exercise to promote skeletal muscle fiber remodeling and myokine IL-15 expression by activating TRPV1 channel": Figure 2-Source Data 1

**Fig. 2D**

Fig. 2D：PGC-1α

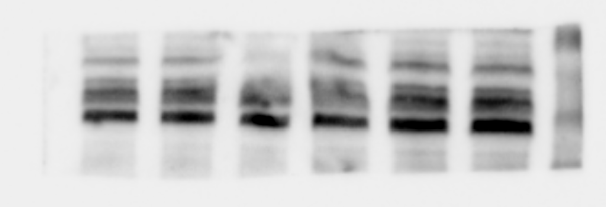

Fig. 2D：Complex V

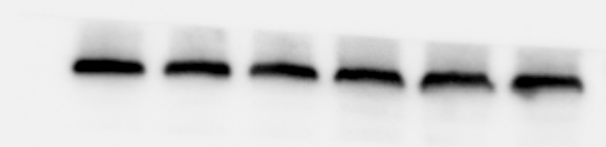

Fig. 2D：Complex IV

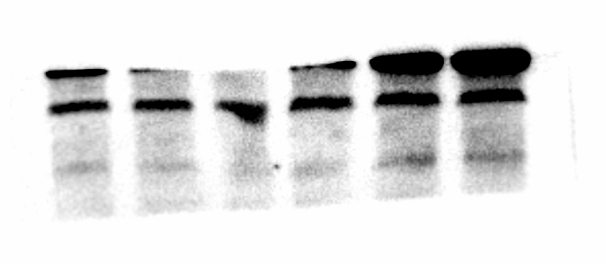

Fig. 2D：Complex III

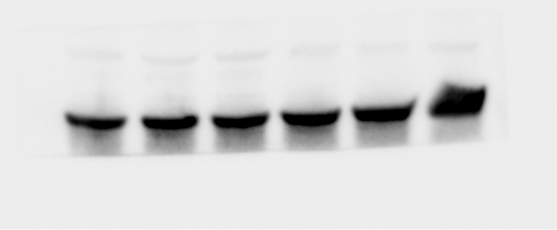

Fig. 2D：Complex II

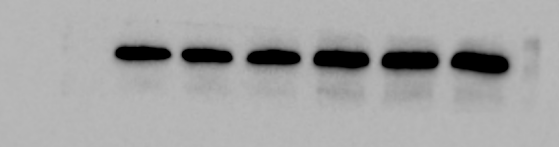

Fig. 2D：Complex I

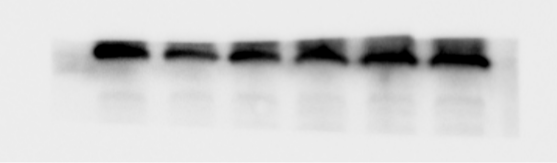

Fig. 2D：β-Actin

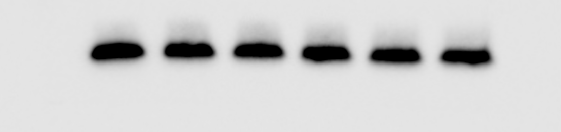
