## Supplementary material for "Eugenol mimics exercise to promote skeletal muscle fiber remodeling and myokine IL-15 expression by activating TRPV1 channel": Figure 3-Source Data 1

**Fig. 3F**

Fig. 3F：iWAT-FABP1

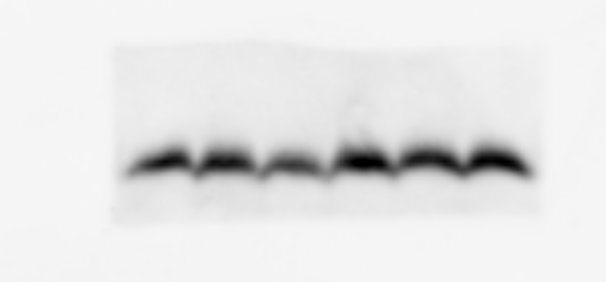

Fig. 3F：iWAT-β-Actin

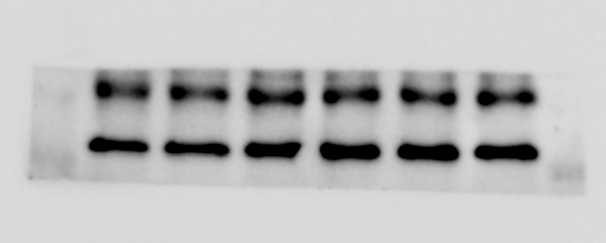

Fig. 3F：gWAT-FABP1

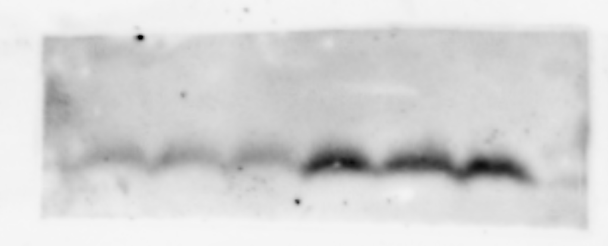

Fig. 3F：gWAT-UCP1

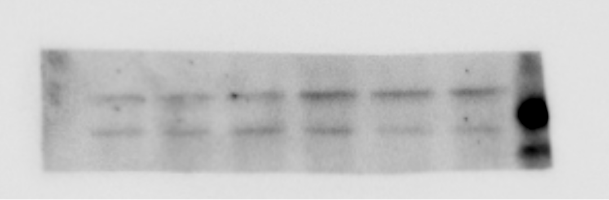

Fig. 3F：gWAT-β-Actin

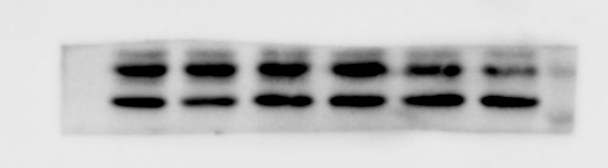

**Fig. 3G**

Fig. 3G：PRDM16

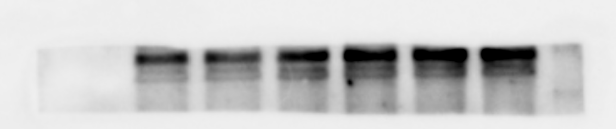

Fig. 3G：UCP1

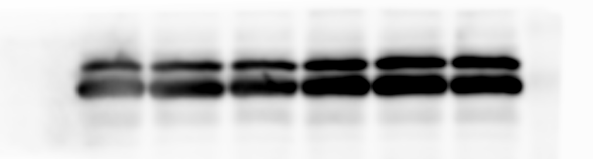

Fig. 3G：PGC-1α

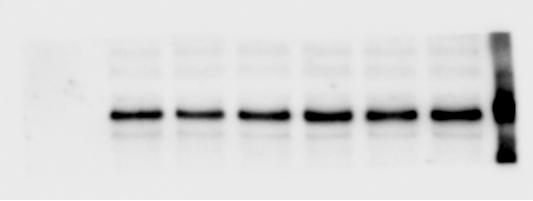

Fig. 3G：

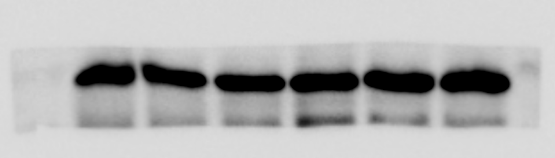

**Fig. 3H**

Fig. 3H：Complex I

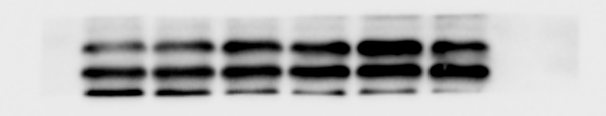

Fig. 3H：Complex II

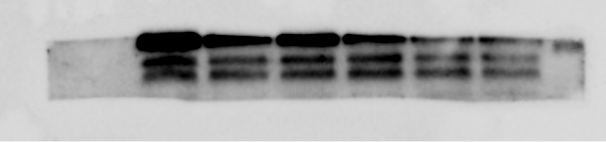

Fig. 3H：Complex III

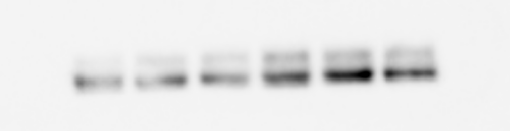

Fig. 3H：Complex IV

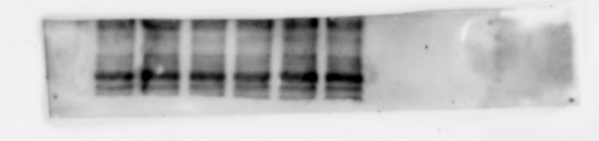

Fig. 3H：Complex V

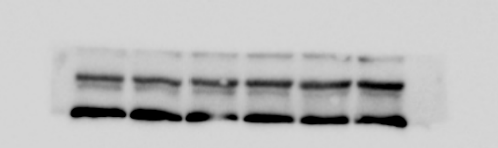

Fig. 3H：β-Actin

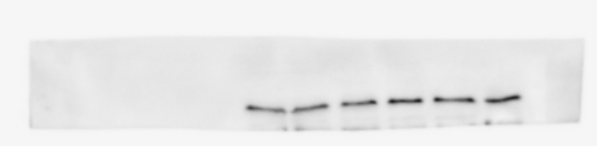
