## Supplementary material for "Eugenol mimics exercise to promote skeletal muscle fiber remodeling and myokine IL-15 expression by activating TRPV1 channel": Figure 4-Source Data 1

**Fig. 4D**

Fig. 4D: TRPV1

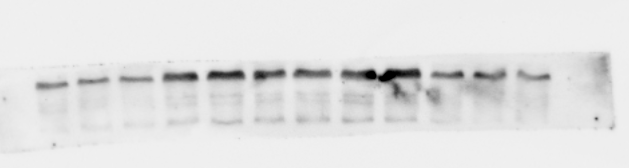

Fig. 4D: CnA

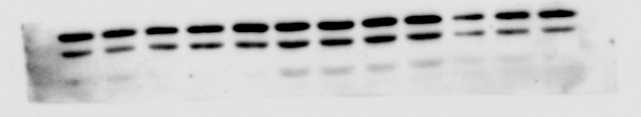

Fig. 4D: β-Actin

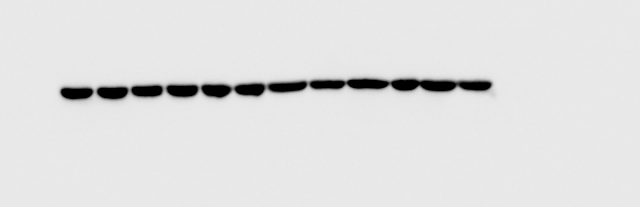

**Fig. 4E**

Fig. 4E: TRPV1

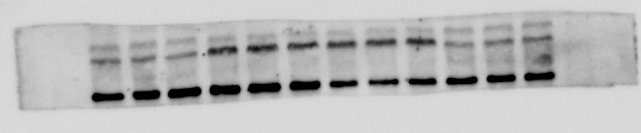

Fig. 4E: CnA

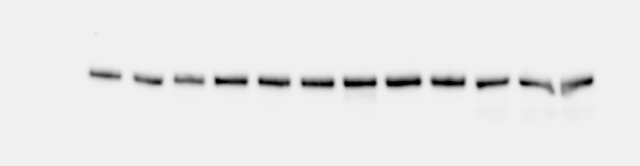

Fig. 4E: β-Actin

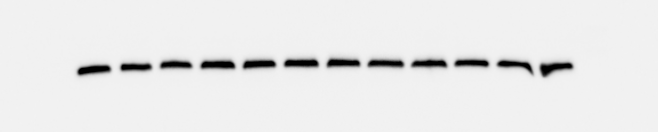

**Fig. 4F**

Fig. 4F: TRPV1

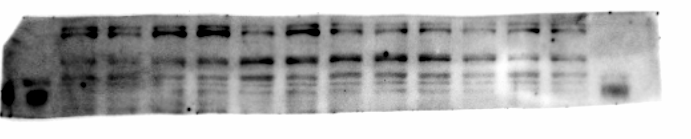

Fig. 4F: CnA

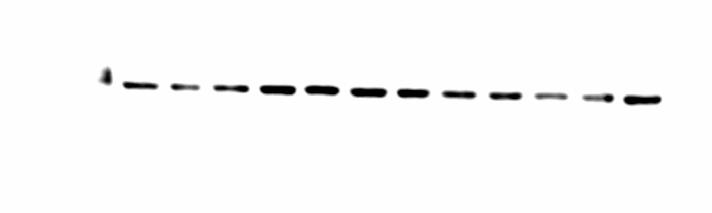

Fig. 4F: β-Actin

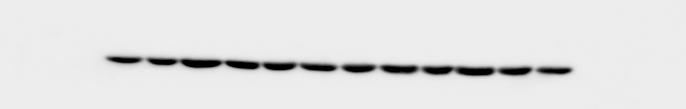

**Fig. 4G**

Fig. 4G: NFATc1

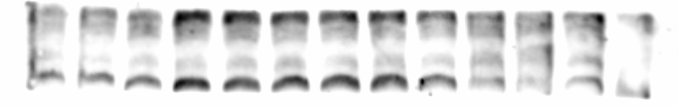

Fig. 4G: Histone H3

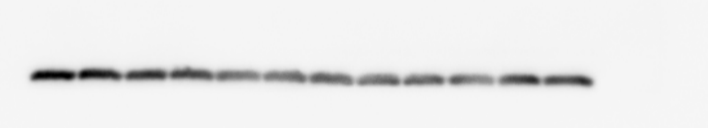

**Fig. 4H**

Fig. 4H: NFATc1

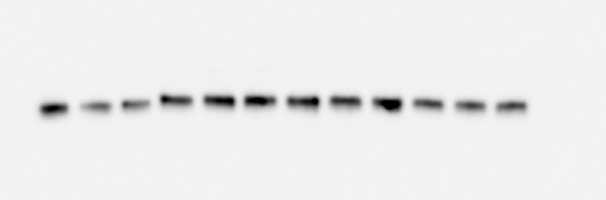

Fig. 4H: Histone H3

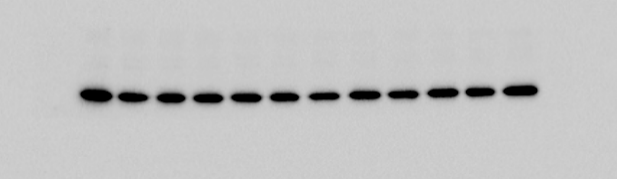

**Fig. 4I**

Fig. 4I: NFATc1

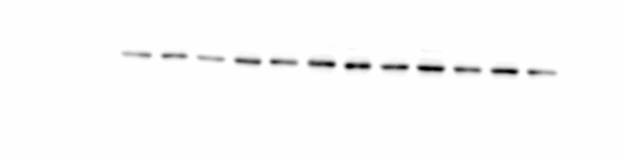

Fig. 4I: Histone H3

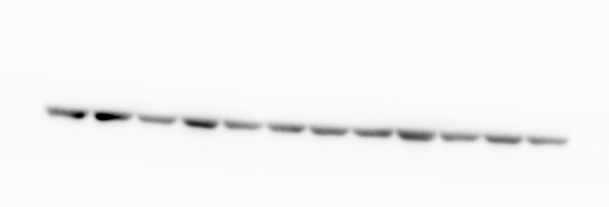
