## Supplementary material for "Eugenol mimics exercise to promote skeletal muscle fiber remodeling and myokine IL-15 expression by activating TRPV1 channel": Figure 5-Source Data 1

**Fig. 5B**

Fig. 5B: CnA

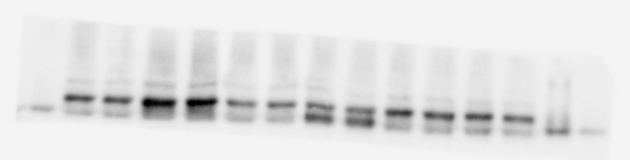

Fig. 5B: β-Actin

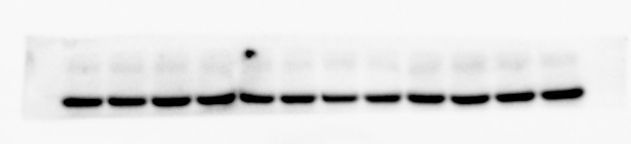

**Fig. 5C**

Figure 5C: Complex I

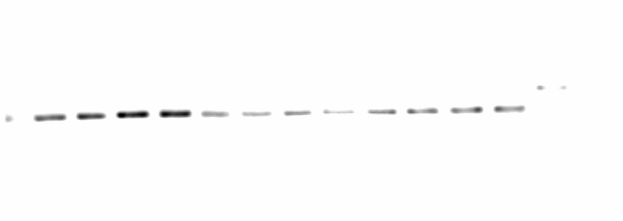

Figure 5C: Complex II

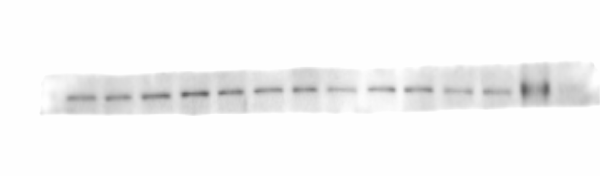

Figure 5C: Complex III

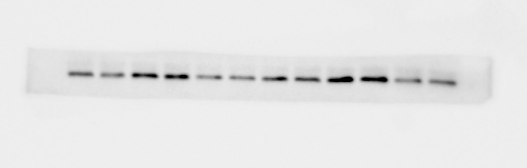

Figure 5C: Complex IV

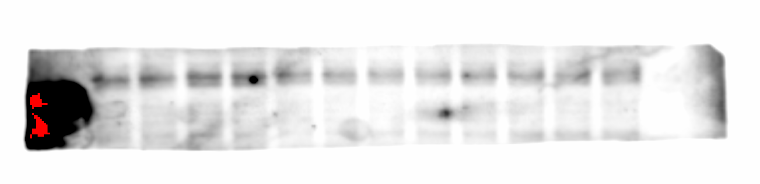

Figure 5C: Complex V

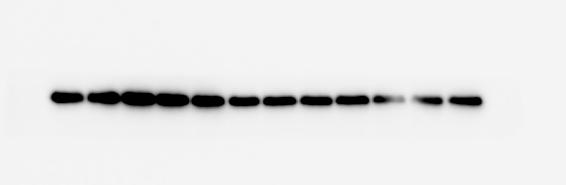

Figure 5C: β-Actin

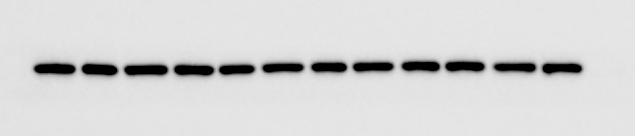

**Fig.5D**

Fig. 5C: Slow MyHC

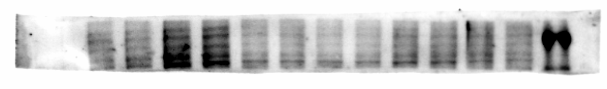

Fig. 5C: Fast MyHC

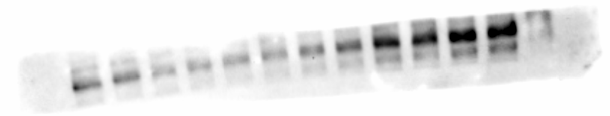

Fig. 5C: β-Actin

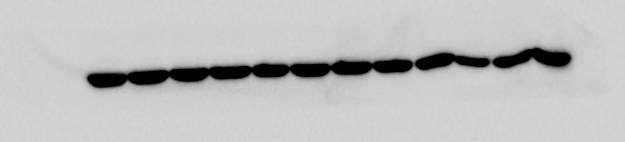
