## Supplementary figures and images for "Eugenol mimics exercise to promote skeletal muscle fiber remodeling and myokine IL-15 expression by activating TRPV1 channel"

### Figure 6-Source Data 1

**Fig. 6B**

Fig. 6B: CnA

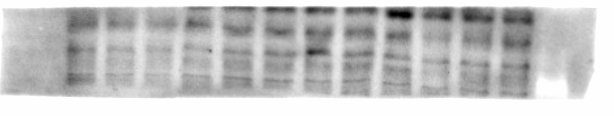

Fig. 6B: β-Actin

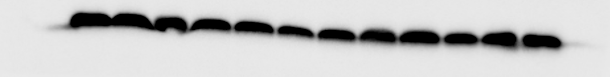

### Figure 9-Source Data 1

**Fig. 9B**

Fig. 9B: IL-15

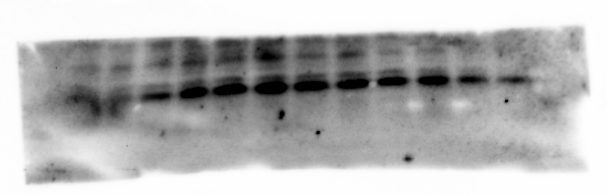

Fig. 9B: Coomassie staining

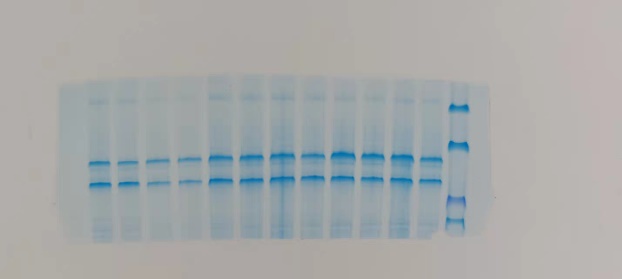

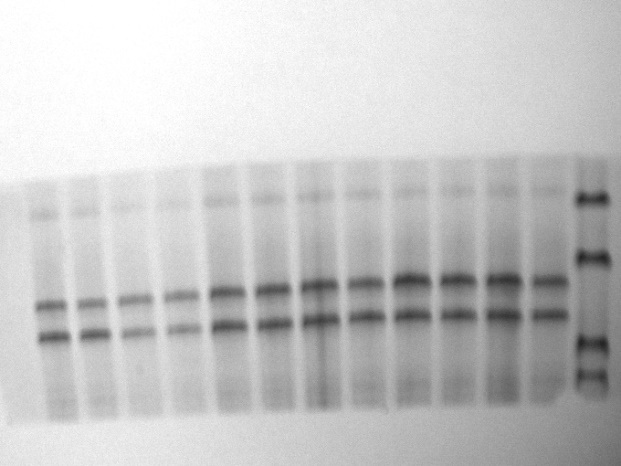

**Fig. 9D**

Fig. 9D: IL-15

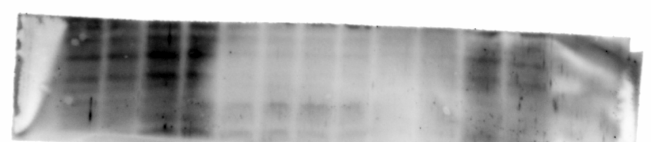

Fig. 8D: β-Actin

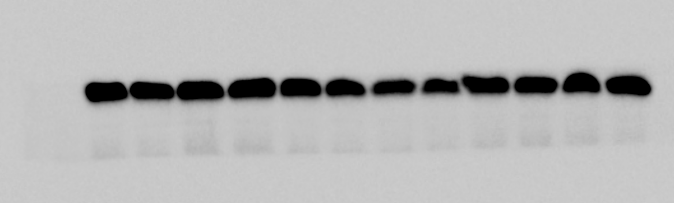
