## Supplementary material for "Eugenol mimics exercise to promote skeletal muscle fiber remodeling and myokine IL-15 expression by activating TRPV1 channel": Figure 7-Source Data 1

**Fig. 7A**

Fig. 7A: IL-15

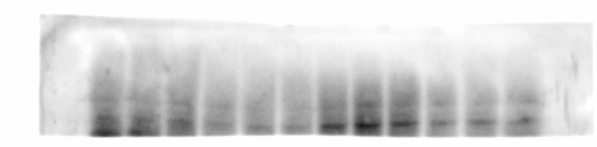

Fig. 7A: β-Actin

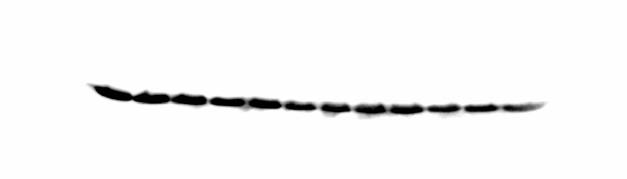

**Fig. 7B**

Fig. 7B: CnA

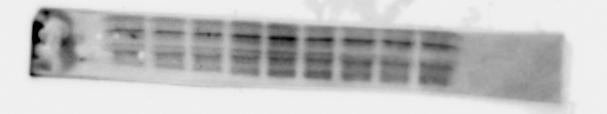

Fig. 7B: β-Actin

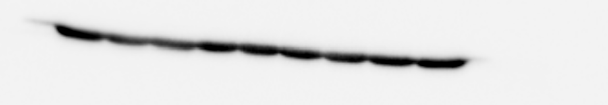

**Fig. 7C**

Fig. 7C: NFATc1

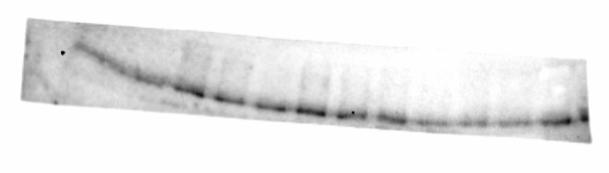

Fig. 7C: Histone H3

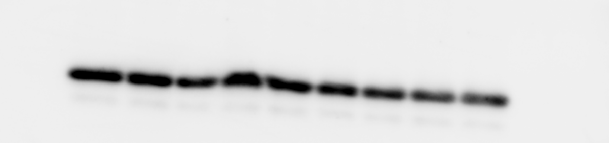

**Fig. 7D**

Fig. 7D: IL-15

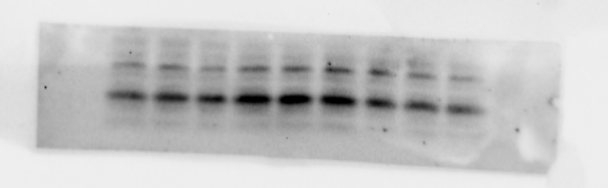

Fig. 7D: β-Actin

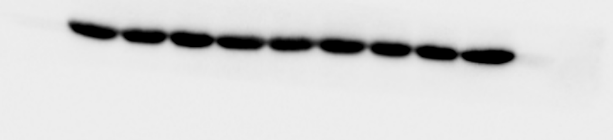

**Fig. 7H**

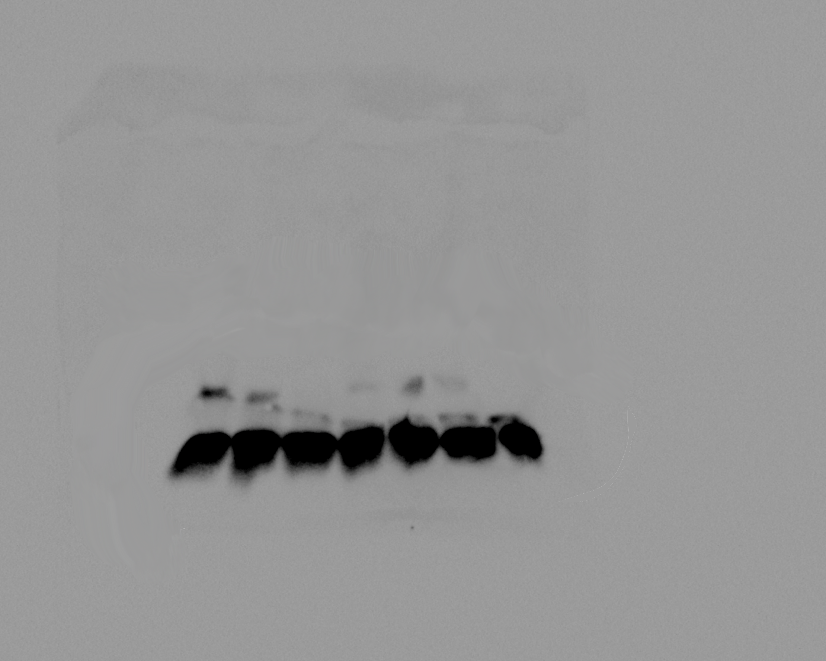

**Fig. 7I**

Fig. 7I: NFATc1

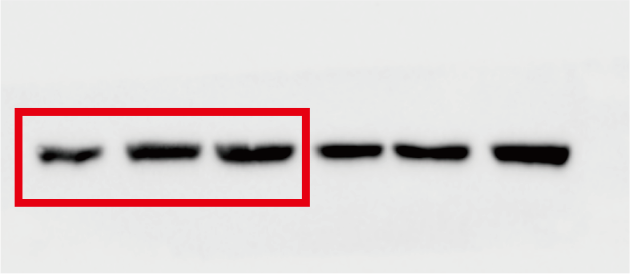

Fig. 7I: β-Actin

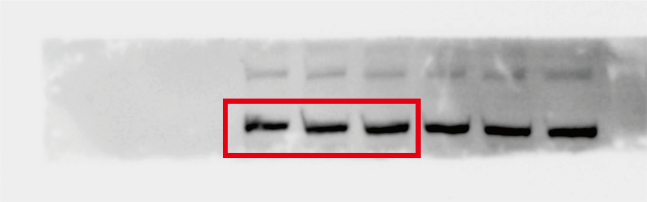
