## Supplementary material for "Eugenol mimics exercise to promote skeletal muscle fiber remodeling and myokine IL-15 expression by activating TRPV1 channel": Figure 8-Source Data 1

**Fig. 8B**

Fig. 8B: IL-15

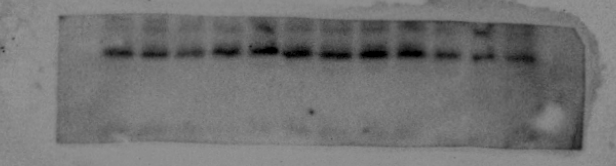

Fig. 8B: β-Actin

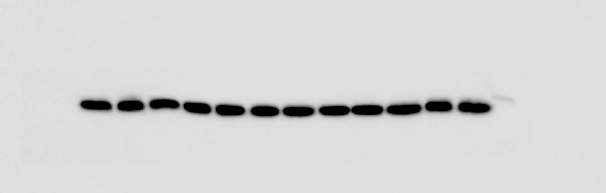

**Fig. 8C**

Fig. 8C: IL-15

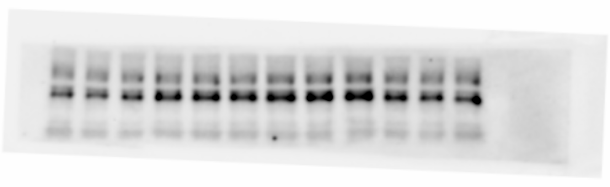

Fig. 8C: β-Actin

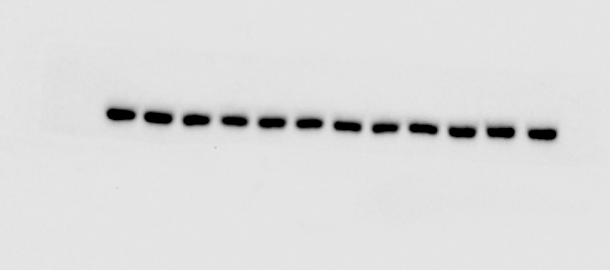

**Fig. 8E**

Fig. 8E: Slow MyHC

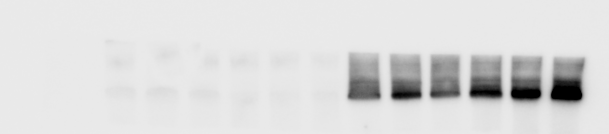

Fig. 8E: Fast MyHC

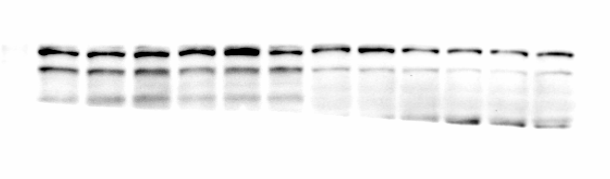

Fig. 8E: IL-15

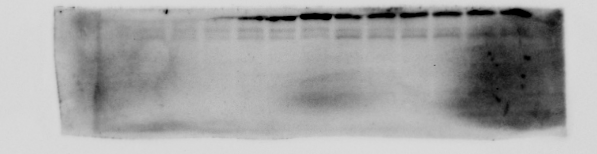

Fig. 8E: β-Actin

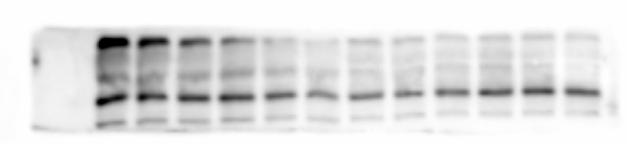
